# RNA triggers chronic stress during neuronal aging

**DOI:** 10.1101/2025.08.04.668575

**Authors:** Kevin Rhine, Elle Epstein, Natasha M. Carlson, Xuezhen Ge, Orel Mizrahi, Anika Kamat, Anita Hermann, William R. Brothers, John Ravits, Eric J. Bennett, Gülçin Pekkurnaz, Gene W. Yeo

## Abstract

Neurodegenerative diseases are linked with dysregulation of the integrated stress response (ISR), which coordinates cellular homeostasis during and after stress events. Cellular stress can arise from several sources, but there is significant disagreement about which stress might contribute to aging and neurodegeneration. Here, we leverage directed transdifferentiation of human fibroblasts into aged neurons to determine the source of ISR activation. We demonstrate that increased accumulation of cytoplasmic double-stranded RNA (dsRNA) activates the eIF2α kinase PKR, which in turn triggers the ISR in aged neurons and leads to sequestration of dsRNA in stress granules. Aged neurons accumulate endogenous mitochondria-derived dsRNA that directly binds to PKR. This mitochondrial dsRNA leaks through damaged mitochondrial membranes and forms cytoplasmic foci in aged neurons. Finally, we demonstrate that PKR inhibition leads to the cessation of stress, resumption of cellular translation, and restoration of RNA-binding protein expression. Together, our results identify a source of RNA stress that destabilizes aged neurons and may contribute to neurodegeneration.

## Introduction

The integrated stress response (ISR) is the cell’s primary defense mechanism against a wide variety of molecular insults, including viral infection, starvation, and the accumulation of unfolded proteins^1^. In stressed conditions, stress responsive kinases will phosphorylate the initiation factor eIF2α, halting the translation of most proteins^2^. The subsequent release of ribosome-associated mRNA leads to the formation of biomolecular condensates called stress granules, which sequester RNA and RNA-binding proteins (RBPs)^3^. Once the stress event has passed, stress granules dissolve and general translation resumes^4^. Overall, the ISR is highly responsive to the cellular environment, activating within minutes of stress exposure and returning to baseline conditions within an hour^4^.

However, dysregulation of the stress response is emerging as a key driver of neurodegeneration^5^. Many neurodegenerative diseases are linked with the formation of pathological aggregates that resemble stress granules^6^; moreover, these pathological aggregates typically consist of RBPs that are released from ribosomes during the canonical ISR^7^. Previous studies have shown that a chronically active ISR in microglia can lead to neurotoxicity in Alzheimer’s disease^8^, and others demonstrated that stress granules may serve a neuroprotective role by preventing runaway aggregation^9^. Preconditioning neurons with small amounts of stress can disrupt the ISR of future stress events, further indicating the fragility of this dynamic cellular process^10^.

We previously demonstrated that neuronal aging leads to chronic activation of the ISR, including basal phosphorylation of eIF2α, altered stress granule RNA binding, and the inability to mount a transcriptional response to new stressors^11^. Aging is one of the primary risk factors for neurodegeneration^12^, so chronic stress in the aging brain may act as a critically important – and therapeutically targetable – initiator of neurodegeneration. However, there is significant disagreement about which ISR pathway might be activated in aging and neurodegenerative diseases, especially because there is evidence of disrupted mitochondrial activity, increased protein aggregation, and faulty RNA metabolism, all of which target distinct ISR kinases^13–16^. Therefore, we sought to identify the source of chronic stress in aged neurons.

Here we show that the accumulation of double-stranded RNA (dsRNA) activates the ISR in aged neurons. Inhibition of the dsRNA-binding kinase PKR, but not other eIF2α kinases, reduces stress granule assembly in aged neurons *in vitro*. We demonstrate that dsRNA levels are elevated in aged neurons cultured *in vitro* and from aged human brain tissue, and this dsRNA sequesters stress granule components in the cytoplasm of aged neurons. The stress-inducing dsRNA is largely mitochondrial, arising from the bidirectional transcription of the mitochondrial chromosome and leaking out from the mitochondria into the cytoplasm to bind PKR. We also show that cellular stress can be reversed by treating transdifferentiated neurons with PKR inhibitor, which also restores translation of RBPs. Together, our results demonstrate the mechanistic basis for the disrupted ISR in aging neurons, providing valuable therapeutic targets that could improve neuronal resiliency by combatting chronic stress.

## Results

### Double-stranded RNA accumulation triggers the formation of stress granules in transdifferentiated neurons

To generate aged human neurons in culture, we transdifferentiated donor fibroblasts directly into neurons (Figure 1A). Transdifferentiation bypasses the pluripotent stem cell state, and transdifferentiated neurons retain key aging markers like altered CpG methylation, expression of senescence and apoptotic markers, and upregulation of aging-associated immune pathways^17–19^. Our transdifferentiation strategy relied on inducible exogenous overexpression of the neurotrophic transcription factors NGN2 and ASCL1 in conjunction with small molecules that antagonized fibroblast cell fates and promoted neuronal maturation (see Methods). Importantly, this differentiation protocol was highly efficient because we used puromycin to eliminate any fibroblasts that did not express NGN2/ASCL1. After three weeks of NGN2/ASCL1 induction, we generated functionally mature neurons that expressed Tubβ3, Map2, and synaptophysin (Figure 1B). RNA-seq analysis confirmed that multiple transdifferentiated lines retained higher expression of senescence and aging-linked markers like *CDKN2A/B* and *IFI16* (Figure S1A-B). In addition, transdifferentiated neurons also expressed immune-linked transcripts at significantly higher levels, including KEGG pathways involved in IL-17 signaling, TNF signaling, and NF-κB signaling (Figure S1C-E, Table S1).

**Figure 1:**
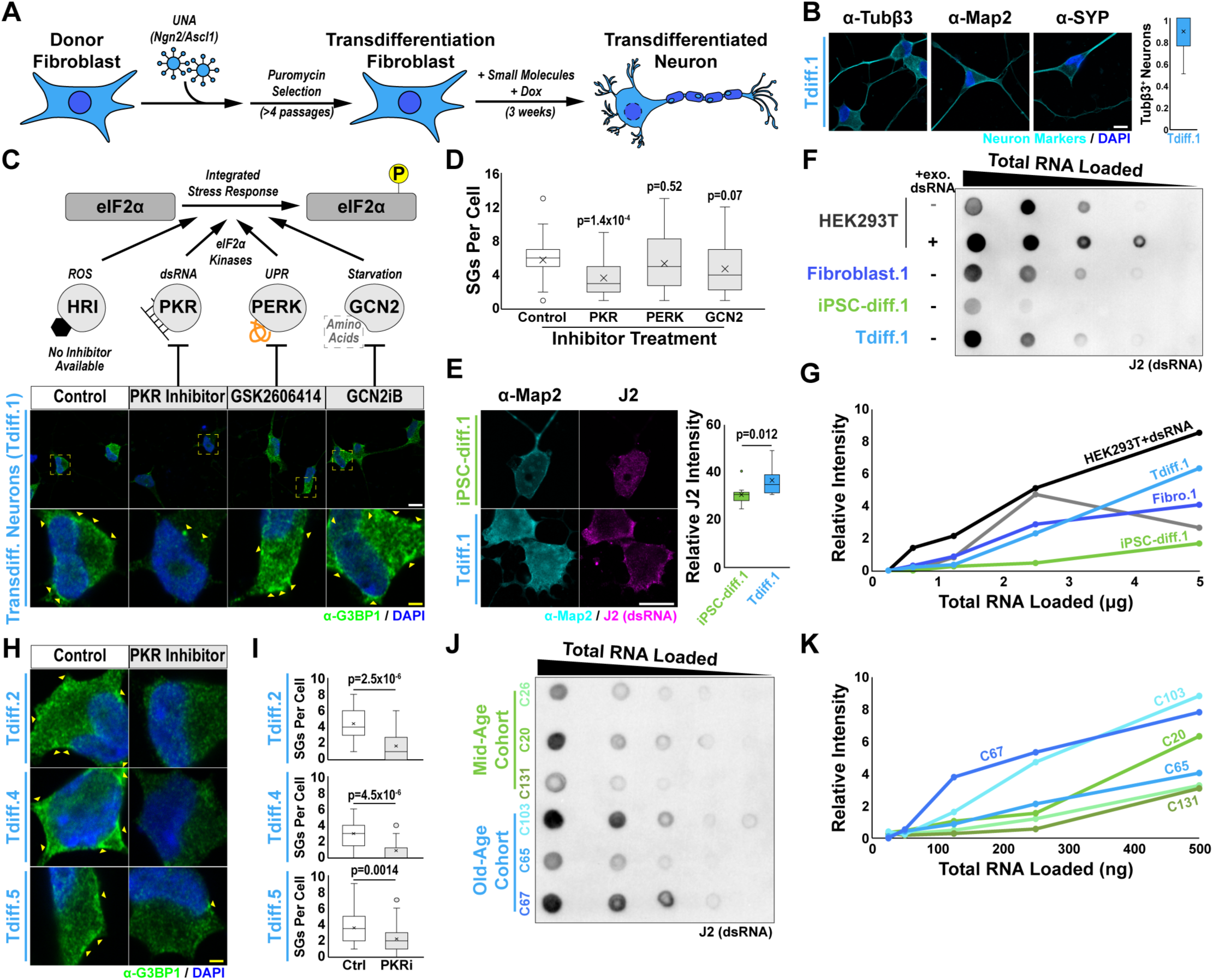
Double-stranded RNA levels are elevated in aged neurons. **(A)** Schematic of neuronal transdifferentiation approach. **(B)** Left, confocal immunofluorescence images of transdifferentiated neurons (Tdiff.1) stained for neuronal markers (cyan) and DAPI (blue). Right, quantification of the fraction of Tubβ3^+^ neurons where the box plot lines denote the 75^th^, 50^th^, and 25^th^ percentile of data values; error bars denote the range of non-outlier values, and the “X” denotes the mean. **(C)** Top, schematic of the eIF2α kinases that activate the integrated response. Bottom, confocal immunofluorescence images of G3BP1 (green) and DAPI (blue) in transdifferentiated neurons (Tdiff.1) treated with the indicated eIF2α kinase inhibitors. The yellow dashed box region denotes the inset, and yellow arrowheads denote chronic stress granules in the cytoplasm of transdifferentiated neurons. White Scale Bar = 10 μm, Yellow Scale Bar = 2 μm. **(D)** Quantification of the number of stress granules (SGs) per cell from the data in (C) (n=3 replicates). From top to bottom, the box plot lines denote the 75^th^, 50^th^, and 25^th^ percentile of data values; error bars denote the range of non-outlier values, the “X” denotes the mean, and any indicated points are outliers. Statistics were calculated using a two-tailed Welch’s t-test for each treatment condition compared to the control. **(E)** Left and middle, representative confocal immunofluorescence images of Map2 (cyan) and dsRNA detected by the J2 antibody (magenta) in transdifferentiated and iPSC-derived neurons. Right, quantification of relative J2 intensity (n=12); the box plot lines denote the 75^th^, 50^th^, and 25^th^ percentile of data values, error bars denote the range of non-outlier values, the “X” denotes the mean, and any indicated points are outliers. White Scale Bar = 10 μm. **(F)** Dot blot of dsRNA intensity detected by J2 antibody staining; total RNA was loaded in decreasing concentrations from left-to-right. Exogenous dsRNA was added to the indicated cells in culture. **(G)** Quantification of dsRNA intensity measured in (F) (n=3 replicates, one example shown). **(H)** Same as in (C) but for the indicated transdifferentiation lines and only treated with the PKR inhibitor. **(I)** Same as (D) but for the data in (H). **(J)** Same as (F) but for RNA isolated from mid-age (green) and old-age (blue) cohorts of brain frontal cortex. **(K)** Same as (G) but for the data in (J).

We previously reported that the stress response was basally activated in transdifferentiated neurons, which led to the formation of chronic stress granules^11^. The integrated stress response is activated by four kinases that respond to different stress stimuli yet all target eIF2α to promote stress granule formation: HRI, which is activated by oxidative stress; PKR, which is activated by double-stranded RNA; PERK, which is activated by the presence of unfolded proteins; and GCN2, which is activated by the loss of essential amino acids (Figure 1C)^13^. To determine which of these kinases might induce stress granule formation in transdifferentiated neurons, we treated the neurons with inhibitors to three of the four kinases (a commercial HRI inhibitor was not available). In our control condition, we observed ∼6 stress granules per neuron as measured by immunofluorescence of G3BP1, a stress granule marker; inhibition of GCN2 and PERK activity did not significantly affect the number of stress granules (Figure 1C-D). However, we observed that PKR inhibition led to a significant 2-fold decrease in the number of stress granules per neuron (Figure 1C-D), indicating that PKR activity was driving stress granule formation in transdifferentiated neurons. PKR inhibition affected the condensation of G3BP1 and eIF2α (Figure S2A), but not the intensity or localization of PKR itself (Figure S2B-C).

Double-stranded RNA (dsRNA) directly binds to PKR, which leads to a conformational change in PKR that enables dimerization followed by autophosphorylation^20,21^. Typically, dsRNA originates from viral infection, but dsRNA can also accumulate from endogenous sources like the mitochondrial transcriptome or dysregulated splicing^22,23^. To determine whether an increase in dsRNA levels might be activating PKR in transdifferentiated neurons, we performed immunofluorescence using the dsRNA-specific J2 antibody, which recognizes dsRNA motifs of >40 bp^24^. Consistent with our inhibitor experiment in Figures 1C-D, we observed higher levels of dsRNA in transdifferentiated neurons relative to isogenic iPSC-derived neurons (Figure 1E). To further confirm that the dsRNA concentration was elevated in transdifferentiated neurons, we performed a dot blot of dsRNA using the J2 antibody. We observed increased dsRNA intensity in isogenic transdifferentiated neurons and fibroblasts – both of which are aged – compared to iPSC-derived neurons and separately in our positive control, HEK293T cells transfected with exogenous dsRNA, relative to untransfected HEK293T cells (Figure 1F-G, Figure S2D-E). Therefore, our results suggest that the accumulation of dsRNA activates PKR and the integrated stress response.

To verify that dsRNA accumulation was not an experimental artifact of the cell line we used (Tdiff.1), we repeated the PKR inhibitor experiment in additional transdifferentiated lines. In three other transdifferentiated neuronal lines (Tdiff.2/4/5), we observed a significant decrease in stress granule formation following PKR inhibitor treatment (Figure 1H-I, Figure S2F-G). We also tested whether dsRNA accumulation was a consequence of transdifferentiation; therefore, we obtained human brain tissue samples from six donors divided into cohorts of mid-aged (∼45 years old) and old-aged (∼80 years old) individuals. We isolated total RNA from the brain samples and repeated the dsRNA dot blot; consistent with our data in transdifferentiated neurons, we observed elevated dsRNA in the old-age cohort relative to the mid-age cohort (Figure 1J-K). Altogether, our data indicate that dsRNA levels increase due to neuronal aging and lead to the activation of PKR.

### Double-stranded RNA binds to stress granule components

The accumulation of dsRNA in transdifferentiated neurons may lead to sequestration of various RBPs, especially stress granule proteins that harbor dsRNA during viral infection^25^ or exogenous dsRNA transfection (Figure S2D). Therefore, we sought to comprehensively identify the proteins interacting with dsRNAs by performing a J2 immunoprecipitation followed by mass spectrometry in Tdiff.1 neurons (Figure 2A). In parallel, we immunoprecipitated dsRNA in isogenic iPSC-derived neurons (iPSC-diff.1), and an IgG immunoprecipitation was used as a control for both J2 pulldowns.

**Figure 2:**
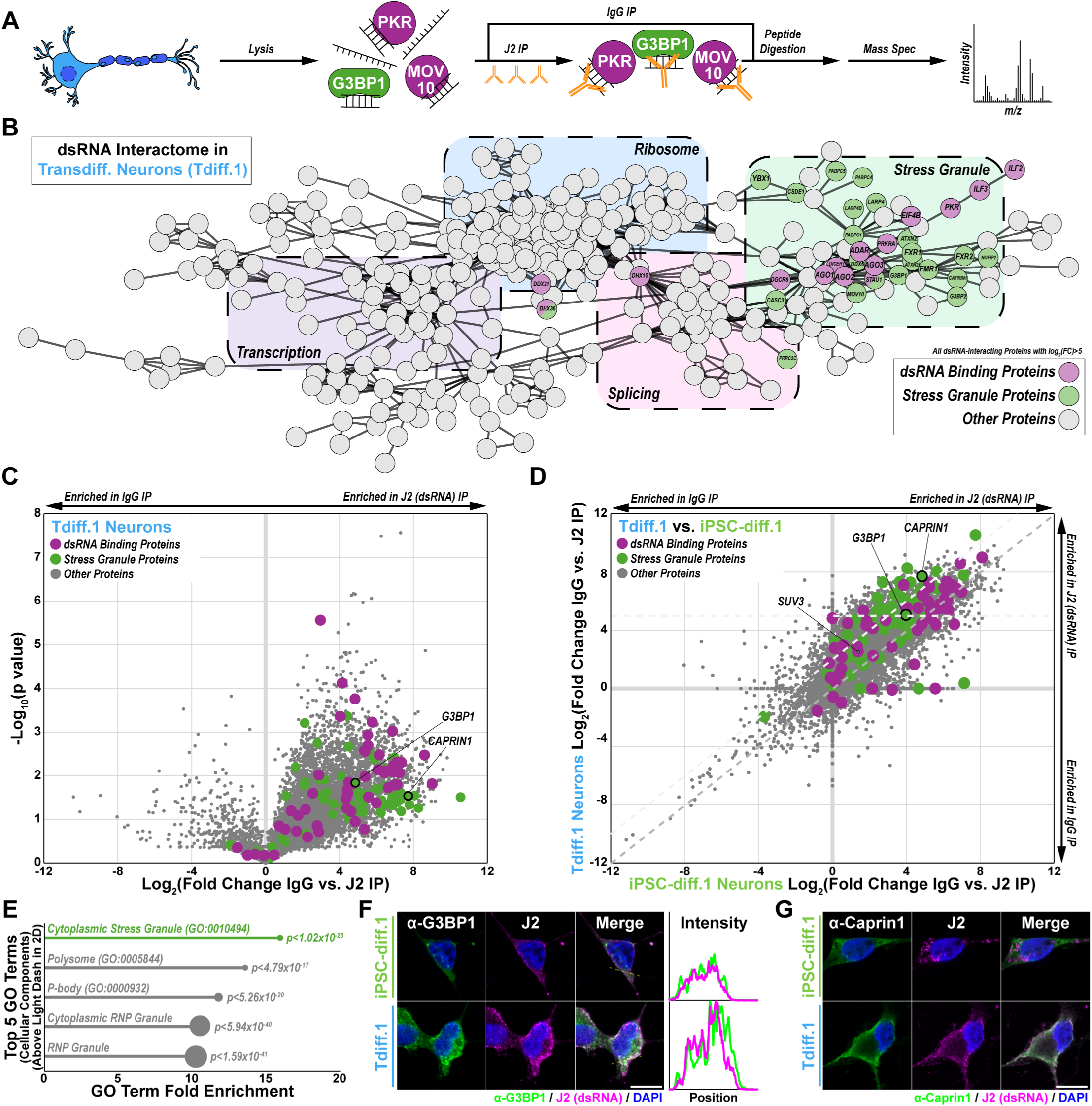
dsRNA sequesters stress granule components in transdifferentiated neurons. **(A)** Schematic of dsRNA (J2) immunoprecipitation and mass spectrometry. **(B)** dbSTRING network of proteins that were immunoprecipitated by dsRNA (J2) in transdifferentiated neurons (Tdiff.1) (n=3 replicates). Only significantly interacting proteins (log_2_(FC)>5 for J2/IgG pulldown) that clustered within the main network are shown. The superposition of distinct groups of proteins (e.g. ribosome, transcription, etc.) is approximate. Nodes of dsRBPs (purple; GO:0003725) and stress granule proteins (green; GO:0010494) are labeled on the network. **(C)** Volcano plot of the log_2_(FC) of dsRNA-interacting proteins in transdifferentiated neurons (Tdiff.1). The p-value was calculated using a two-tailed Welch’s t-test. Proteins of interest are labeled as in (B), and the canonical stress granule markers G3BP1 and Caprin1 are labeled. **(D)** Fold change versus fold change plot of J2/IgG pulldown in transdifferentiated neurons and isogenic iPSC-derived neurons. dsRBPs and stress granule proteins are highlighted as in (B). The gray dashed line denotes the y=x line whereas the light gray dashed lines denote log_2_(FC)>5 for the Tdiff.1 pulldown and log_2_(FC-Tdiff.1) −log_2_(FC-iPSC.diff.1)>2. **(E)** Lollipop plot of the top 5 Gene Ontology terms for the proteins above the light gray dashed lines in (D); the size of the circle scales with the number of proteins within each term. **(F)** Co-immunofluorescence and intensity profile of G3BP1 and J2 (dsRNA) in the indicated neurons. White Scale Bar = 10 μm. **(G)** Same as (F) but for Caprin1.

We found that more than 800 proteins were enriched (log_2_(Fold Change(J2/IgG))>5) in the J2 immuno-precipitation of transdifferentiated neurons (Figure 2B). These proteins were largely RBPs, encompassing diverse portions of the RNA life cycle like translation, transcription, and splicing (Figure 2B, Table S2). As expected, double-stranded RNA binding was a top gene ontology term (Figure S3A, Table S2); many well-known dsRBPs were immunoprecipitated, including ADAR, AGO1/AGO2/AGO3, PKR, DICER1, and STAU1 (Figure 2B)^26–28^. Importantly, the cytoplasmic stress granule was also a top gene ontology term (Figure S3B, Table S2), and the interactome of dsRNA-immunoprecipitated stress granule RBPs was highly interconnected with dsRBPs (Figure 2B).

A few stress granule proteins were previously reported as weak dsRNA-interacting proteins^27,29^, but we hypothesized that aging might further enrich stress granule proteins relative to unaged neurons. Indeed, we found that canonical stress granule proteins like G3BP1/2, Caprin1, ATXN2/2L, and the FMR/FXR family were highly enriched in the dsRNA interactome of transdifferentiated neurons but not iPSC-derived neurons (Figure 2B-C, Figure S3C-D, Table S3). When the enrichment of all dsRNA-interacting proteins in transdifferentiated neurons was mapped to the enrichment in isogenic iPSC-derived neurons, stress granule proteins emerged as the top gene ontology term for proteins that were more highly enriched in the transdifferentiated neurons (Figure 3D-E, Table S3). P-body proteins were also stronger dsRNA interactors in transdifferentiated neurons (Figure 3E), indicating that there may be widespread accumulation of dsRNA in cytoplasmic RNA granules. By contrast, other RNA pathways that made up a large portion of our network in Figure 2B (e.g. ribosomal proteins) were not over-enriched in the transdifferentiated neurons (Figure S3E), indicating that the sequestration of stress granule RBPs by dsRNA due to aging is likely specific to their function in the stress response.

**Figure 3:**
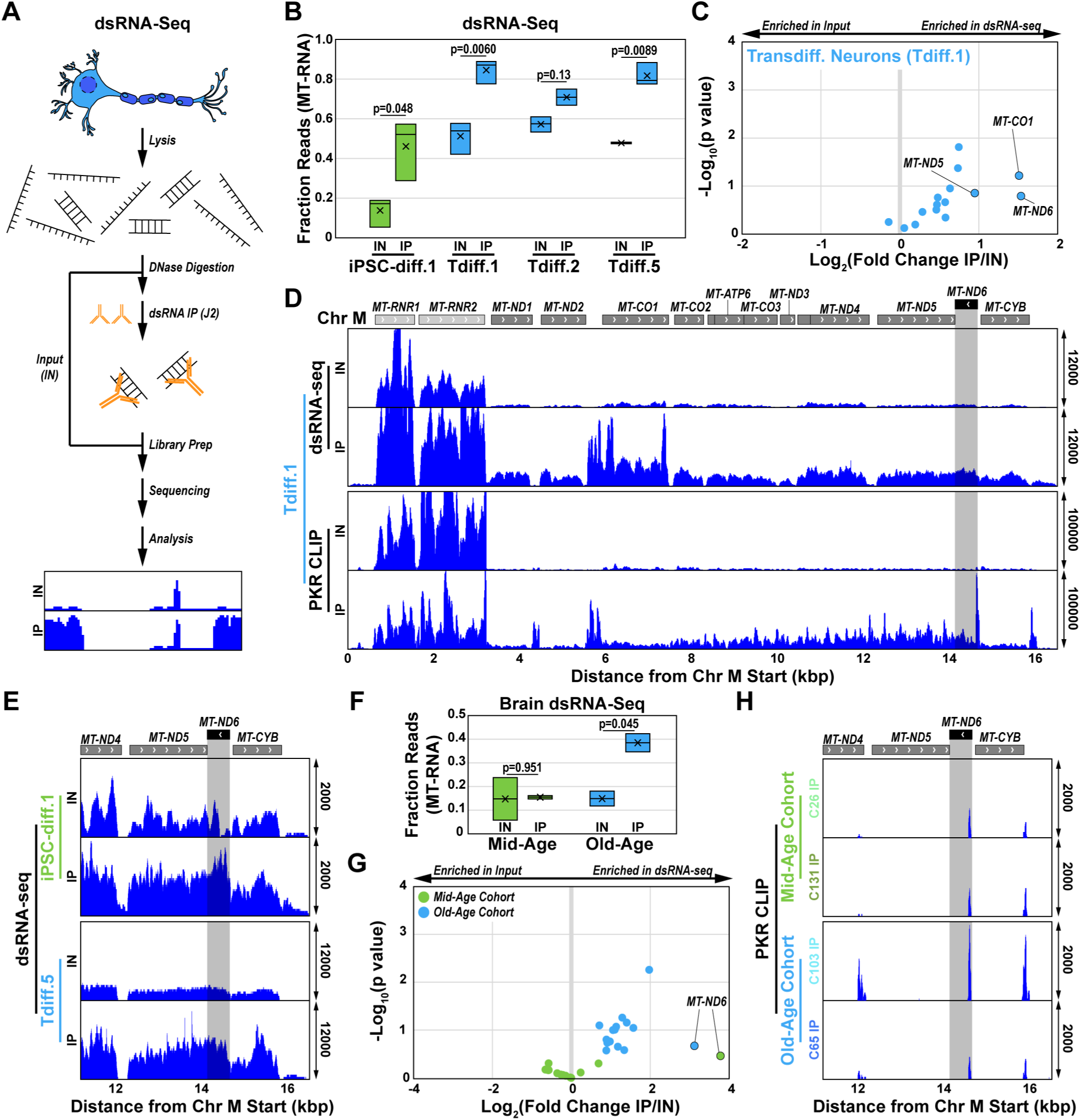
Aging leads to accumulation of mitochondrial dsRNA in neurons. **(A)** Schematic of the dsRNA-seq protocol. **(B)** Boxplot of the fraction of transcripts identified as mitochondrial (MT-)RNA in the input (IN) and immunoprecipitation (IP) samples of the indicated cell lines (n=3 replicates). **(C)** Volcano plot of the log_2_(Fold Change) of all MT-RNA transcripts identified by dsRNA-Seq in the input and IP samples of the Tdiff.1 transdifferentiated neuron line. **(D)** Visualization of representative mitochondrial sequencing reads detected by dsRNA-Seq (n=3 replicates) and PKR CLIP (n=2 replicates) in the Tdiff.1 transdifferentiated neuron line. The shaded region denotes the reverse-transcribed *MT-ND6* gene. The range of the y-axis values is denoted on the right side of the genomic track and was fixed for each pair of input and IP samples. **(E)** Same as (D) but for the ∼11-17 kb inset of the mitochondrial chromosome for the dsRNA-sequencing results from the indicated cell lines. **(F)** Same as (B) but for cohorts of human brain tissue (N=2). **(G)** Same as (C) but for cohorts of human brain tissue (N=2). **(H)** Same as (D) but for the indicated human brain tissue samples.

To confirm that stress granule components were bound by dsRNA in transdifferentiated neurons, we performed immunofluorescence of two key stress granule RBPs from our network: G3BP1 and Caprin1 (Figure 2B-C). Consistent with our mass spectrometry results, we observed that dsRNA and stress granule proteins colocalized into bright foci in transdifferentiated neurons (Figure 2F-G). By contrast, G3BP1, Caprin1, and dsRNA remained diffuse in iPSC-derived neurons (Figure 2F-G), again demonstrating that there is an age-dependent interaction between stress granules and dsRNA.

### Endogenous mitochondrial dsRNA activates the stress response in aged neurons

Although dsRNA accumulation is typically a consequence of exogenous introduction via a viral infection, we hypothesized that endogenous dsRNA might be accumulating due to aging. Endogenous dsRNA can form in response to bidirectional transcription of the nuclear or mitochondrial genome, loss of circular RNAs, genotoxic stress, or decreased autophagy^23^. Therefore, we performed dsRNA-sequencing, which relies on J2-mediated immunoprecipitation of dsRNA (Figure 3A)^30^. We confirmed that this approach successfully recovered dsRNA in HEK293T cells by tracking exogenously transfected dsRNA throughout the purification and immunoprecipitation protocol (Figure S4A). To further validate our recovery of dsRNA, we demonstrated that the dsRNA-specific exonuclease RNase III degraded our immunoprecipitated dsRNA (Figure S4A).

Next, we performed dsRNA-sequencing on transdifferentiated neurons. We surprisingly found that more than 80% of the immunoprecipitated dsRNA was derived from mitochondrial transcripts in three distinct transdifferentiated neuronal lines (Figure 3B, Table S4). Previous studies have also reported that the majority of endogenous dsRNA is derived from mitochondrial genome^30–32^, which is bidirectionally transcribed as two long polycistronic transcripts^33^. The sole mitochondrial protein-coding RNA that is transcribed in the reverse orientation, MT-ND6, was the most enriched transcript in dsRNA pulldowns relative to the input samples (Figure 3C-E, Figure S4B-C). Most other mitochondrial mRNAs – which are transcribed in the forward orientation – were also enriched in the immunoprecipitated dsRNA compared to the corresponding input samples (Figure 3C-E, Figure S4B-C). We also observed enrichment of mitochondrial dsRNA in iPSC-derived neurons (Figure 3B, Figure S4D, Table S4), indicating that the mitochondrial transcriptome is also the predominant source of dsRNA in young neurons. Therefore, we sought to determine whether the accumulation of mitochondrial dsRNA is uniquely stressful to aged neurons. We revisited our bulk RNA-seq data and analyzed the expression of nuclear-encoded mitochondrial transcripts (GO:0005739) and mitochondrial transcripts (MT-RNA); we found that MT-RNA was expressed significantly higher compared to all genes and to nuclear-encoded mitochondrial transcripts in transdifferentiated neurons relative to iPSC-derived neurons (Figure S4E-F). In addition, the nuclear-encoded genes for *SUPV3L1* (encoding SUV3) and *PNPT1* – which localize to the mitochondria to unwind and degrade MT-dsRNA, respectively – were depleted in transdifferentiated neurons (Figure S4E).

As mentioned above, PKR binds to dsRNA to initiate the ISR. To further confirm that MT-dsRNA was activating the integrated stress response via PKR, we performed enhanced crosslinking and immunoprecipitation (eCLIP)-sequencing of PKR-bound RNA in transdifferentiated and iPSC-derived neurons^34,35^. In transdifferentiated neurons, we observed PKR binding at specific loci in the mitochondrial chromosome in Tdiff.1 neurons: the 5’ end of the MT-ND6 gene, the intergenic region between MT-ND1 and MT-ND2, and the intergenic region between MT-ND2/MT-CO1 (Figure 3D). By contrast, these peaks were weak relative to input in iPSC-derived neurons, again demonstrating that the MT-dsRNA phenotypes are unique to aged neurons (Figure S4F).

To confirm that mitochondrial dsRNA also increased in the aging human brain, we repeated the dsRNA-seq and PKR eCLIP-seq experiments in postmortem brain tissue. Consistent with our results in transdifferentiated neurons, we observed strong enrichment of MT-dsRNA in the old-age cohort (Figure 3F-G, Table S5); the enrichment of MT-dsRNA was considerably weaker in the mid-age cohort (Figure 3F-G, Table S5). In addition, there was a strong PKR binding peak at the MT-ND6 transcript like in transdifferentiated neurons (Figure 3H). Together, our results demonstrate that (1) the vast majority of dsRNA in aged neurons is from the mitochondrial transcriptome, (2) this MT-dsRNA directly binds to PKR to activate the ISR, (3) MT-dsRNA in iPSC-derived neurons is not sufficient to elicit PKR activation, and (4) the human brain accumulates MT-dsRNA as a function of aging.

### Damaged mitochondria leak stress-inducing dsRNA into the cytoplasm of transdifferentiated neurons

Mitochondrial damage was previously reported to increase in both aged neurons and in the neurons of patients with neurodegenerative diseases^36,37^; MT-dsRNA was also previously shown to activate the innate immune response via MDA5 and MAVS in senescent cells^31,32^. Therefore, we hypothesized that mitochondrial damage might lead to leakage of MT-dsRNA into the cytoplasm, which would then elicit the PKR-mediated stress response that we observed. To test our hypothesis, we determined whether MT-ND6 – the most enriched transcript in our dsRNA-seq and PKR CLIP results in Figure 3 – was localized outside of the mitochondria.

We performed immunofluorescence of the mitochondrial marker TOM70 coupled with fluorescence *in situ* hybridization (FISH) of the MT-ND6 transcript. Consistent with our hypothesis, we found MT-ND6 foci that were localized outside of the TOM70 signal in four transdifferentiated neuron lines (Tdiff.1/2/4/5), including in synapses and in the soma (Figure 4A, Figure S5A). Meanwhile, the mitochondrial ribosomal RNA (RNR1) colocalized completely with TOM70 (Figure S5B), which aligns with our data showing that RNR1 was not a major target of PKR (Figure 3D). Other mitochondrial markers like PDH, COXIV, and TOM20 were also constrained to the mitochondria (Figure S5C).

**Figure 4:**
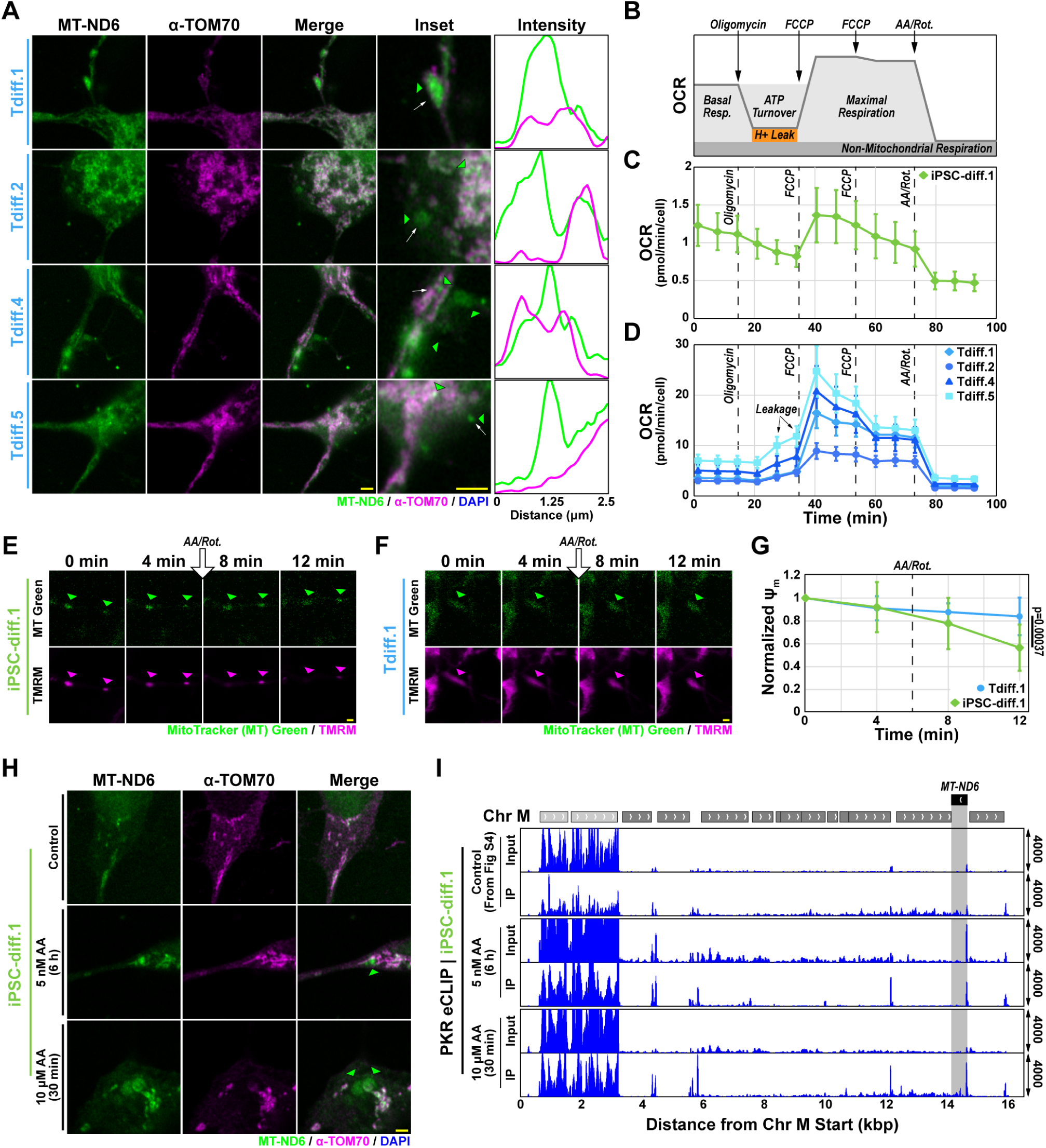
Leaky mitochondria release mitochondrial dsRNA. **(A)** Airyscan immunofluorescence-FISH images of the mitochondrial marker TOM70 (magenta) and MT-ND6 transcript (green) in transdifferentiated neurons (Tdiff.1/2/4/5). Green arrowheads denote cytoplasmic MT-ND6 puncta; the white arrows denote the intensity profiles in the adjacent plot. Yellow Scale Bar = 2 μm. **(B)** Schematic of typical mitochondrial stress test results. **(C)** Plot of oxygen consumption rate (OCR) in the mitochondrial stress test (1 μM of each compound) of iPSC-diff.1 neurons (n=6 replicates). Error bars are standard deviation. **(D)** Same as (C) but for Tdiff.1-5 neurons with 3 μM of each compound. **(E)** Fluorescence confocal microscopy images of representative mitochondria in the neurites of iPSC-diff.1 neurons. The mitochondria were visualized with TMRM (magenta) and MitoTracker (MT) Green (green) and are identified with arrowheads. Rotenone (Rot.) and antimycin A (AA) were added 6 min into the video as indicated by the white arrow. Yellow Scale Bar = 2 μm. **(F)** Same as (E) but for Tdiff.1 neurons. **(G)** Quantification of ψ_m_ as a function of time in the perfusion experiments in panels (E) and (F) (n=3 replicates). Error bars denote standard deviation, and statistics were calculated using Welch’s t-test. **(H)** Same as (A) but for iPSC-diff.1 treated with DMSO (control), 5 nM AA for 6 h, or 10 μM AA for 30 min (n=3 replicates). The green arrowheads denote cytoplasmic MT-ND6 puncta. **(I)** Representative genomic tracks of input and immunoprecipitation (IP) samples of PKR CLIP in iPSC-diff.1 neurons treated with AA as in panel (G) (n=2 replicates). The PKR CLIP track for the control condition is reproduced from Figure S4G. The shaded region denotes the MT-ND6 gene.

Mitochondrial dsRNA is processed by the helicase SUV3 and degraded by the exonuclease PNPT1^31^, so we wondered whether these proteins might be malfunctioning in aged neurons. Although we found that these proteins were mostly localized mostly in the mitochondria and still bound to the mitochondrial dsRNA as measured by eCLIP (Figure S5C-E), there were also several experiments that indicated a failure to properly process MT-dsRNA. First, the binding profile of SUV3 included scattered peaks throughout the mitochondrial chromosome in transdifferentiated neurons but not iPSC-derived neurons (Figure S5E). Second, our dsRNA AP-MS experiment from Figure 2D also showed stronger SUV3 binding to dsRNA in transdifferentiated neurons than in iPSC-derived neurons. And third, SUV3 and PNPT1 were depleted in transdifferentiated neurons despite a significantly higher load of MT-RNA overall and MT-dsRNA (Figure 3B, Figure S4E). Therefore, our results suggest that SUV3 and PNPT1 are not able to properly process the toxically high loads of MT-dsRNA.

We next sought to determine how MT-ND6 and other MT-dsRNAs might escape from mitochondria. To elucidate the status of the mitochondrial membrane, we performed a mitochondrial stress test and measured the oxygen consumption rate (OCR) in Tdiff.1/2/4/5 and iPSC-diff.1 neurons^38^. In iPSC-derived neurons, we observed the canonical OCR changes in the mitochondrial stress test: OCR attenuation in response to oligomycin treatment, strong OCR stimulation in response to carbonyl cyanide 4-(trifluoromethoxy)phenylhydrazone (FCCP), and cessation of all respiration in response to antimycin A and rotenone (Figure 4B-C). By contrast, the transdifferentiated neuronal lines all failed to maintain OCR attenuation following oligomycin treatment (Figure 4D). Because oligomycin inhibits ATP synthesis and proton translocation, we reasoned that increased doses of oligomycin might restore the canonical suppression of proton movement. However, high concentrations of oligomycin (up to 5 μM) also failed to suppress proton leakage (Figure S6A-B), indicating that this result was not due to improper dosage. Instead, our results suggest that protons steadily leak across the mitochondrial membrane in aged neurons^39^, which might also enable dsRNA to leak if the membrane is sufficiently damaged^32^.

Therefore, we performed live-cell imaging of Tdiff.1 mitochondria to determine whether they were damaged at baseline. The mitochondrial membrane potential (ψ_m_) – which is maintained by an intact mitochondrial membrane^40^ to support ATP production – can be approximated by tetramethylrhodamine (TMRM)^41,42^. We stained Tdiff.1 and the isogenic iPSC-diff.1 neurons with TMRM and normalized the signal to MitoTracker Green, which is not affected by changes in ψ_m_ (Figure S6C-D). Consistent with our hypothesis, we found that ψ_m_ of Tdiff.1 mitochondria were insensitive to antimycin A and rotenone treatment (Figure 4E-G). Together, our data in Tdiff.1 neurons show that their mitochondria is highly dysfunctional and likely damaged enough to support dsRNA leakiness into the cytoplasm.

To further mechanistically link mitochondrial leakiness to the PKR-mediated stress response, we treated iPSC-diff.1 neurons with antimycin A, an antibiotic that induces mitochondrial damage and depolarization. We used two different regimes of antimycin A treatment: a long incubation with a low concentration (5 nM for 6 h) and a short incubation with a high concentration (10 μM for 30 min)^43^. IF-FISH showed that both conditions led to accumulation of cytoplasmic MT-ND6 foci that were distinct from TOM70-positive mitochondria (Figure 4H). In addition, we observed stronger PKR binding to MT-dsRNA, especially to MT-ND6 in the 10 μM antimycin A condition (Figure 4I). Together, our results demonstrate that aging-linked mitochondrial damage leads to MT-dsRNA release, which then activates the stress response via PKR.

### PKR inhibition in aged neurons restores ribosomal function

Finally, we determined the extent to which PKR inhibition might restore normal cellular functions in transdifferentiated neurons, especially the resumption of translation. Therefore, we treated Tdiff.1 neurons with PKR inhibitor and performed several stress- and translation-focused experiments, including bulk RNA-seq, phospho-eIF2α blotting, G3BP1 immunoprecipitation and mass spectrometry, and Riboseq (Figure 5A).

**Figure 5:**
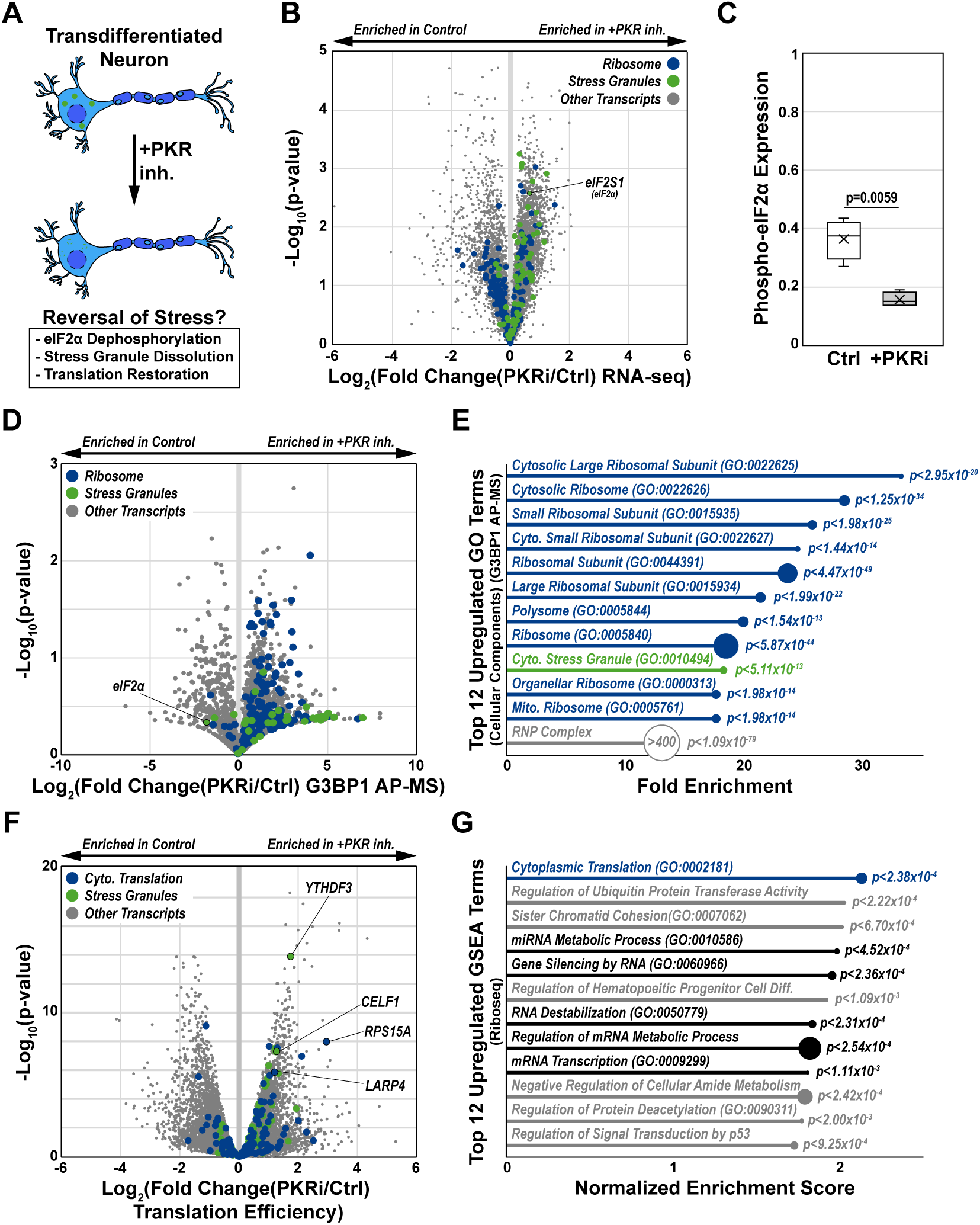
PKR inhibition reverses cellular stress and restores ribosomal function. **(A)** Schematic of the PKR inhibitor experiments in this figure. **(B)** Volcano plot of RNA-seq expression of all detected transcripts in transdifferentiated neurons (Tdiff.1) with or without PKR inhibitor treatment (n=3 replicates). The p-values were calculated using a two-tailed Welch’s t-test. Stress granule transcripts are highlighted in green (GO:0010494) and ribosomal transcripts are highlighted in blue (GO:0005840). **(C)** Quantification of phospho-eIF2α expression relative to total eIF2α expression in transdifferentiated neurons (Tdiff.1) with and without PKR inhibitor treatment as detected by Western blot (n=4 replicates). Statistics were calculated using a two-tailed Welch’s t-test. **(D)** Volcano plot of proteins detected by G3BP1 AP-MS in transdifferentiated neurons (Tdiff.1) with or without PKR inhibitor treatment (n=3 replicates). Stress granule proteins are highlighted in green (GO:0010494) and ribosomal proteins are highlighted in blue (GO:0005840). P-values were calculated using a two-tailed Welch’s t-test. **(E)** Lollipop plot of top gene ontology terms calculated by ShinyGO v0.82 among proteins that were enriched >4-fold in the G3BP1 AP-MS data from panel D. The size of the circle correlates to the number of genes in the corresponding term. Ribosomal pathways are highlighted in blue, and the stress granule pathway is highlighted in green. **(F)** Volcano plot of mRNA transcripts detected by Riboseq in Tdiff.1 neurons with or without PKR inhibitor treatment (n=2 replicates). Stress granule proteins are highlighted in green (GO:0010494) and cytoplasmic translation proteins are highlighted in blue (GO:0002181). P-values were calculated using a two-tailed Welch’s t-test. **(G)** Lollipop of normalized enrichment scores of top gene ontology terms from data in (F). The size of the circle correlates to the number of genes in the corresponding term. Ribosomal pathways are highlighted in blue, RBP metabolic pathways are highlighted in black.

In our previous study, we found that RBPs were depleted due to neuronal aging^11^. Here, PKR inhibition led to increased expression of transcripts encoding predicted RBPs (Figure S7A-B)^44^. Interestingly, some classes of RBPs (e.g. stress granule components) were upregulated whereas other groups of RBPs (e.g. the ribosome) were depleted (Figure 5B, Figure S7B). The ISR marker eIF2S1 (encoding eIF2α), which is both a ribosomal protein and a stress granule component, was significantly enriched after PKR inhibition (Figure 5B). Because phosphorylated eIF2α is the canonical marker for ISR activation, we assayed whether phospho-eIF2α levels would decrease after PKR inhibitor treatment in Tdiff.1 neurons. Indeed, we found that PKR inhibition indeed led to a significant reduction in phospho-eIF2α levels (Figure 5C, Figure S7C), again indicating a reduction in stress levels.

Phosphorylation of eIF2α leads to stalled translation where key stress granule components leave the ribosome and assemble into condensates. Therefore, we tested whether PKR inhibition also impacted the protein-protein interaction network of a key stress granule protein, G3BP1. Immunoprecipitation of G3BP1 and mass spectrometry of the interacting proteins revealed a striking restoration of G3BP1-ribosomal interactions (Figure 5D-E, Table S6). This result is consistent with previous literature showing that G3BP1 stabilizes the ribosome and promotes translation in unstressed conditions^45,46^. Unlike other ribosomal proteins, eIF2α was less enriched in the G3BP1 interactome of PKR inhibitor-treated neurons, likely reflecting dissolution of the stress granule. We also observed stronger G3BP1 binding to other stress granule components like Caprin1 (Figure 5D-E); however, many of these proteins are also linked to translation in steady-state conditions^47^.

To test whether this restored G3BP1 interaction with the ribosome might also lead to functional changes in translation, we performed Riboseq in PKR-inhibitor treated and control Tdiff.1 neurons^48^. By calculating the translation efficiency of each transcript from bulk RNA-seq and Ribo-seq, we found that PKR inhibition led to increased translation of transcripts encoding diverse RNA metabolic pathways, including translation, stress granules, and RNA degradation (Figure 5F-G, Figure S7D-E). Overall, this led to increased translation efficiency of predicted RBPs (Figure S7D-E), which may help restore RNA biology in aged neurons.

## Discussion

Mitochondrial dysfunction is strongly associated with neurodegenerative diseases^49^; here, we find that neuronal aging leads to the accumulation of dsRNA from the mitochondrial genome, which triggers the chronic stress response. In both transdifferentiated neurons and the aging human brain, we demonstrate that there are significantly higher levels of MT-dsRNA that directly bind to PKR and sequester stress granule components. PKR inhibition leads to a broad reversal of stress phenotypes that plague aged neurons, especially the resumption of general translation and the dissolution of stress granules. However, this PKR-focused intervention does not address the cause of MT-dsRNA leakage: aging mitochondria that fail to properly produce energy for the cell, are unable to properly seal their membranes, and cannot degrade accumulating MT-dsRNA. Together, our results recast a known culprit in neurodegeneration – the damaged mitochondria – in a new light by delineating an RNA pathway that chronically exacerbates cellular stress in aged neurons (Figure 6).

**Figure 6:**
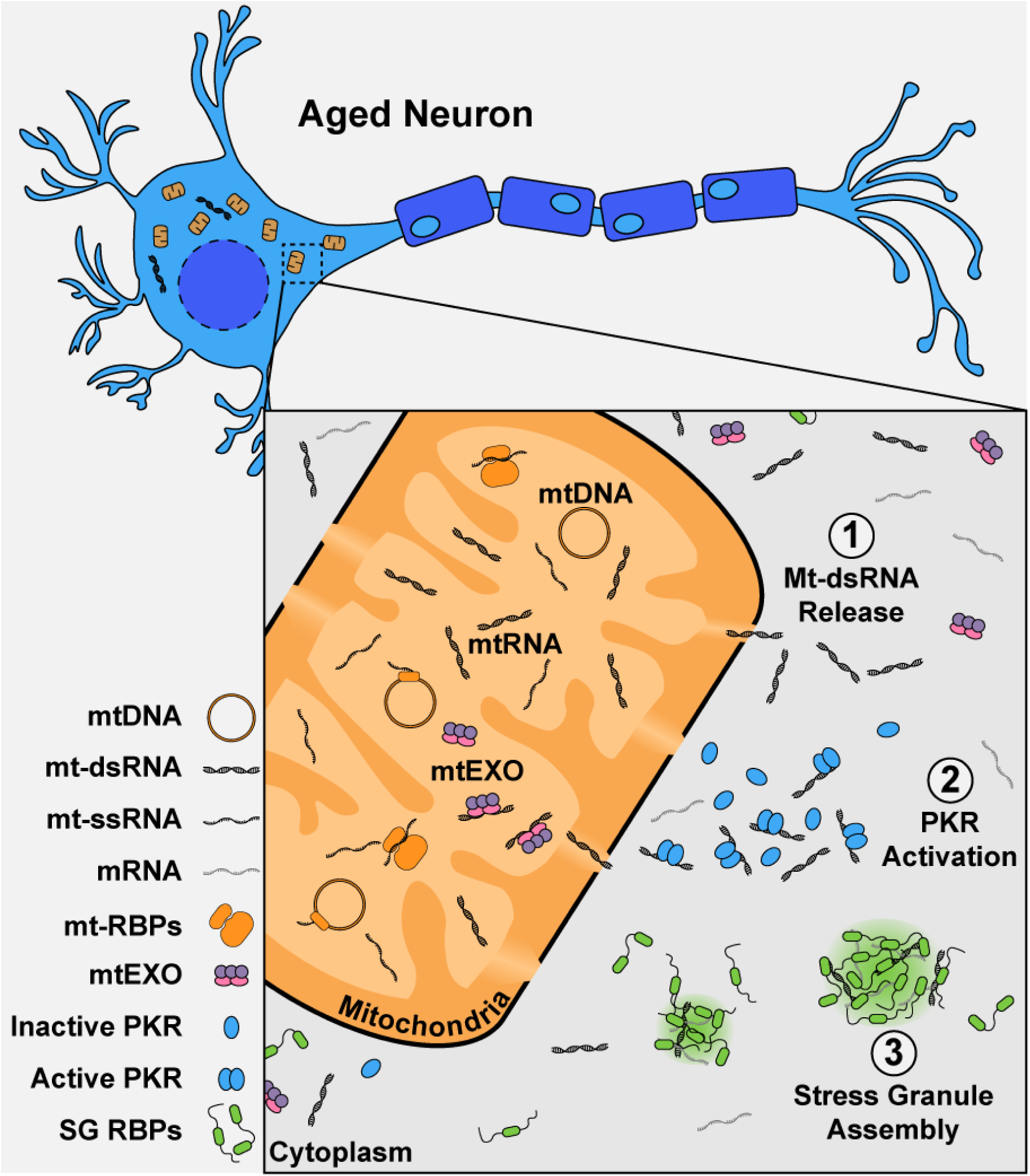
Leaky mitochondria cause chronic activation of the stress response in aged neurons. Our results indicate that neuronal aging leads to (1) MT-dsRNA release, (2) PKR activation, and (3) stress granule assembly. The chronic stress response can be ablated in aged neurons with PKR inhibitor or induced in young neurons with Antimycin A.

Transdifferentiation is emerging as a unique and tractable approach to study neuronal aging. Previously, our group and others have shown that transdifferentiated neurons retain aging markers yet also mature into functional neurons that can form synapses and fire action potentials^11,18,50,51^. Our results likewise demonstrate that transdifferentiation retains hallmarks of aging like the overexpression of immune transcripts and senescence markers (Figure S1). The unique benefit of transdifferentiated neurons is that they can be manipulated *in vitro* to study human aging^19^; in this study, we treat transdifferentiated neurons with small-molecule inhibitors, pause translation with cycloheximide for Riboseq, and perform live-cell tracking of mitochondria. These types of experiments are not possible in postmortem human brain tissue, and other model organisms do not fully recapitulate human biology^19^. Transdifferentiation enables us to first identify novel results *in vitro* to design targeted experiments in postmortem human brain tissue; for example, we show that dsRNA levels increase due to aging alongside a concurrent increase in MT-dsRNA load first in transdifferentiated neurons and then in postmortem human brain tissue (Figure 1J-K, Figure 3F-G). Our use of transdifferentiated neurons coupled with key experiments in postmortem human brain tissue enables us to carefully and mechanistically dissect the underpinnings of aging-linked stress in neurons.

It is likely that many stress pathways are simultaneously activated in neurodegeneration, and previous studies have demonstrated the contributions of distinct signaling cascades. In particular, unfolded proteins accumulate in both aged and degenerating neurons, which bind PERK to initiate eIF2α phosphorylation^52–54^. Other ISR kinases like GCN2 may also contribute to neurodegeneration by responding to metabolic deficiencies^55,56^, and indeed GCN2 inhibition was nearly significant in reducing stress granule formation in our results (Figure 1C-D). The mitochondrion itself likely directly or indirectly triggers PERK, GCN2, and even other pathways like cGAS/STING via MT-DNA release^57–60^. However, the importance of PKR follows a plethora of recent literature highlighting the immunogenic potential of dysregulated RNA^22,23,61–64^ and the role of innate immunity in exacerbating neurodegeneration^65–67^.

Endogenous dsRNA is tightly regulated to ensure that it does not activate PKR like exogenous dsRNA from a viral infection^25,61^. In the nucleus, repetitive elements are repressed to prevent the transcription of these dsRNA-prone sequences and splicing is tightly regulated to prevent aberrant backsplicing^68,69^. In the cytoplasm, RNAs are coated with RBPs that inhibit dsRNA hybridization or sensing through RNA modifications, competitive binding, and exonuclease activity^61,70–72^. However, mitochondria are also a major source of dsRNA: neurons typically have hundreds of mitochondria each with several copies of MT-DNA that generate long bidirectional dsRNA^73,74^. This MT-dsRNA is peeled apart by SUV3 helicase and degraded by the exonuclease PNPT1^31^. In aged neurons, the incredibly high load of MT-dsRNA likely overwhelms these quality control mechanisms, leading to the enriched MT-RNA detected in both bulk and dsRNA-sequencing (Figure 3B, Figure S4E). This dsRNA can then interact with PKR^22^, as we show here, or with other dsRNA sensors like MAVS/MDA5 and RIG-I^32^.

Our results demonstrate that a concurrent loss of mitochondrial membrane integrity offers an opportunity for MT-dsRNA to escape into the cytoplasm and bind to PKR. We observed direct evidence of MT-ND6 RNA outside mitochondria in transdifferentiated neurons; the same effect could be artificially induced in iPSC-derived neurons by treating with antimycin A (Figure 4H-I). However, it is unclear how the mitochondrial membrane integrity is lost in the aging brain. A previous study showed that senescent cells had mitochondria with visible protrusions that led to dsRNA release^32^. The cell should clear these damaged mitochondria through mitophagy, but the intense energy needs of a neuron and the high load of other misfolded proteins may prevent prompt turnover of mitochondria^59,75^. Another study suggested that PKR coats, and may even enter, the mitochondria to bind to MT-dsRNA^22^, but we do not observe evidence of intramitochondrial PKR via immunofluorescence or in our control PKR CLIP experiments (Figure S2C, S4G). Future studies should focus on the mechanism of how mitochondrial homeostasis is lost throughout neuronal aging, contributing to PKR activation.

The failure to resolve MT-dsRNA chronically triggers the ISR and leads to the formation of stress granules. These stress granules sequester dsRNA as detected by mass spectrometry (Figure 2B-D). In addition, translation is stalled as G3BP1 and other stress granule proteins leave the ribosome; this effect can be reversed by treating with PKR inhibitor. Although PKR inhibitor can reverse the deleterious effects of MT-dsRNA on an aged neuron, it is likely not a viable therapeutic because innate immune pathways cannot be wholly deactivated. A more targeted therapeutic that directly unwinds or degrades MT-dsRNA may be more clinically feasible. In light of our previous work^11^, such a therapeutic could restore neuronal resiliency to new stress and help prevent the onset of neurodegenerative diseases like amyotrophic lateral sclerosis and Alzheimer’s disease. Together, our results highlight the importance of RNA regulation in aging, neuronal homeostasis, and neurodegeneration: RNA acts as both the initiator of stress and the effector of compromised neuronal function.

## Acknowledgements

We thank Katherine Rothamel for helpful discussions and other Yeo lab members (Brian Yee, Grady Nguyen, Steven Blue, and Sara Mumford) for technical support; we also thank Zach Whiddon for assistance with live-cell mitochondrial imaging. Sequencing, imaging, and proteomics support was provided by the Institute for Genomic Medicine core staff at UCSD, the Advanced Biophotonics core staff at the Salk Institute, and the Proteomics core staff at Sanford Burnham Prebys, respectively. We acknowledge funding support from the following sources: the Milton Safenowitz Postdoctoral Fellowship (23-PDF-639) and a Seed Grant (26-SGP-764) from the ALS Association to K.R; the UCSD Eureka! Scholars Program to E.E.; the NIH (T32-EB009380) to N.M.C.; the Gruss Lipper Postdoctoral Research Fellowship by the EGL Charitable Foundation to O.M.; the UCSD Ledell Family Research Scholarship for Science and Engineering to A.K.; FightMND, Target ALS, and the NIH (R01-NS069566) to A.H. and J.R.; Target ALS (IL-2023-C2-L4) and the NIH (R35-GM148339) to X.G. and E.J.B.; the NIH (R35-GM128823) to G.P.; and Grant 2023-332369 from the Chan Zuckerberg Initiative DAF, an advised fund of Silicon Valley Community Foundation, and the NIH (R01-HG004659 and R01-NS103172) to G.W.Y.

## Contributions

K.R. and G.W.Y. conceived of the project. K.R. performed cell culture, inhibitor treatment assays, bulk RNA sequencing, dsRNA-seq library preparation, eCLIP-Seq, affinity pulldowns, immunofluorescence, and IF-FISH experiments, prepared statistical analyses for the paper, coordinated the project, and wrote the primary draft of the manuscript; E.E. performed dsRNA dot blots, exogenous dsRNA synthesis, dsRNA-seq immunoprecipitation and assisted with cell culture and imaging data analysis; N.C. performed and analyzed the data from mitochondrial stress tests and live-cell imaging of mitochondrial membrane potential; X.G. performed the dsRNA immunoprecipitation mass spectrometry and processed mass spectrometry raw data; O.M. performed and analyzed Riboseq; A.K. assisted with mitochondrial immunofluorescence/IF-FISH experiments and cell culture; A.H. curated human brain samples; W.R.B. assisted with ribosomal experiments; J.R. provided fibroblast lines and human brain tissue; E.J.B. supervised dsRNA mass spectrometry experiments; G.P. supervised mitochondrial imaging and respirometry experiments; G.W.Y supervised the project and edited the final manuscript. All authors contributed to the final version of the manuscript.

## Data and Materials Availability

Raw and processed data are available at the following biorepositories: all RNA-sequencing data (including bulk, dsRNA-sequencing, Ribo-seq, and eCLIP-seq) at NIH GEO with the accession code GSE301408; dsRNA immunoprecipitation and mass spectrometry at PRIDE with the accession code PXD066264; G3BP1 immunoprecipitation and mass spectrometry at PRIDE with the accession code PXD066084; and raw imaging data at Mendeley Data (doi:https://10.17632/s25558n7cp.1).

## Code Availability Statement

No new code or software was created for this manuscript.

## Methods

### HEK-293T / Lenti-X 293T Culturing

HEK-293T (ATCC #CRL-3216) and Lenti-X 293T (Takara #632180) cells were maintained in DMEM (high glucose, Life #11965118) supplemented with 10% (v/v) fetal bovine serum (Life #26140079) at 37 °C with 5% CO_2_. For passaging, 90+% confluent cells were treated with TrypLE Express (Life #12604013) for 5 min and split in the desired ratio with fresh DMEM + 10% FBS.

### Lentiviral Production

Lenti-X 293T cells were passaged to a fresh plate at ∼50% confluency. Within 24 h of a fresh passage, the newly-seeded cells were co-transfected with psPAX2 (Addgene #12260), pMD2.G (Addgene #12259), and pLVX-UbC-rtTA-Ngn2:2A:Ascl1 (UNA; Addgene #127289) or pLVX-UbC-rtTA-Ngn2:2A:EGFP (rtTA-Ngn2; Addgene #127288) at a 1:1:2 mass ratio with the JetPEI transfection system (VWR #89129-916). The supernatant fraction of the transfected cells was collected 48 h, 72 h, and 96 h and pooled at 4 °C. The collected virus was supplemented with Lenti-X concentrator (Takara #631232), incubated at 4 °C for 30 min, and centrifuged at 1500 x *g* for 45 min at 4 °C. Following centrifugation, the lentivirus-containing pellet was resuspended in 1X dPBS (Life #13190250), the viral titer was measured using Lenti Go-Stix (Takara #631280), and the purified virus was diluted to the desired titer for storage at -80 °C.

### Fibroblast Cell Culture & Transduction

In brief, fibroblasts were cultured in TFM (DMEM, high glucose; 15% (v/v) fetal bovine serum; 1X MEM NEAA, Thermo #11140050) at 37 °C with 5% CO_2_. TFM was replaced every 2-3 days, and fibroblasts were always maintained at >30% confluency. For passaging, fibroblasts were detached with TrypLE Express for 5 min and split at 1:2 or 1:3 split ratios. Long-term fibroblast stocks were resuspended in BamBanker (Bulldog Bio #BB05) for storage in liquid nitrogen. To generate transdifferentiation-competent (UNA) fibroblasts, 80% confluent cells were seeded on 6-well plates (Fisher #FB012927) incubated in 500 μL TFM with lentivirus containing the UNA expression vector in 5 μg/mL polybrene (Sigma #TR-1003-G) for 8 h at 37 °C. Afterwards, the transduction volume was increased to 2 mL for an additional 36 h incubation. The transduced fibroblasts were then incubated in TFM-P (TFM with 1 μg/mL puromycin, Thermo #A1113803) for at least 4 passages to select for transdifferentiation-competent fibroblasts. Cultures with death rates greater than 50% after selection were discarded.

### Fibroblast Transdifferentiation

Plates for transdifferentiation were coated with 10 μg/mL poly-D-lysine (PDL; Sigma #P6407-5MG) and 0.0001% (w/v) poly-L-ornithine (PLO; Sigma #P4957-50ML) for 24 h and 10 μg/mL laminin (Sigma #L2020-1MG) for an additional 24 h. The coated plates were seeded with 300% confluent UNA fibroblasts. Transdifferentiating neurons were cultured for three weeks in NK media (NK) (0.5X DMEM/F-12, GlutaMAX supplement, Life #10565042; 0.5X Neurobasal, Life #21103049; 1X B-27 supplement, Life #17504044; 1X N-2 supplement, Life #17502048; 20 μg/mL laminin; 400 μg/mL dibutyryl-cAMP (db-cAMP), SelleckChem #S7858; 2 μg/L doxycycline, Sigma #D9881-10G; 5 μM dorsomorphine homolog 1 (DMH1), Fisher #412610; 0.5 μM LDN-193189, Fisher #50176042; 0.5 μM A83-01, Stem Cell Technologies #72022; 5 μM forskolin, Stem Cell Technologies #72114; 3 μM CHIR-99021, Fisher #442310; 10 μM SB-431542, Fisher #161410; 10 U/mL penicillin-streptomycin, Thermo #15140122) with media changes every 48-72 h. The transdifferentiation progress was checked every 48-72 h using a Zeiss Axio-Vert.A1. The transdifferentiated neurons used in this study are listed in Table S7.

### iPSC Neuron Differentiation

Previously reprogrammed induced pluripotent stem cells (iPSCs) were maintained in mTeSR Plus (Stem Cell Technologies #100-0276) on plates coated with 1X Matrigel (Corning #354277). The mTeSR media was changed every 48 h. For differentiation, iPSCs were transduced with the rtTA-Ngn2 lentivirus in mTeSR Plus without polybrene. Puromycin (1 μg/mL) was added 48 h after lentiviral transduction. The puromycin-resistant iPSCs were used for neuronal differentiation; these cells were seeded at ∼25% confluency in mTeSR Plus with 2 μg/mL doxycycline for 72 h at 37 °C with 5% CO_2_. After 72 h, the cells were transitioned to NMM1 media (0.5X DMEM/F12, GlutaMAX supplement; 0.5X Neurobasal; 1X B-27 supplement; 1X N-2 supplement; 2 μg/mL laminin; 0.5 mM db-cAMP; 20 ng/μL recombinant BDNF, Peprotech #450-02-50UG; 20 ng/μL human GDNF, Peprotech #450-10-50UG; 10 ng/μL human NT-3, Peprotech #450-03-50UG; 10 U/mL penicillin-streptomycin) with half-media changes every 48 h. Doxycyline was removed after seven days of differentiation; the cells were maintained in NMM1 without doxycycline for an additional two weeks to ensure complete differentiation.

### Small-Molecule and Inhibitor Treatments

Cells were treated as indicated with the following inhibitors: PKR inhibitor (Sigma #527450) at 1 μM for 24 h; GCN2iB (MedChemExpress #HY-112654) at 5 μM for 24 h; GSK2606414 (MedChemExpress #HY-18072) at 1 μM for 24 h; antimycin A (Sigma #A8674) at 5 nM for 6 h or 10 μM for 30 min, except for the Mitochondrial Stress Test as described below.

### dsRNA Synthesis and Transfection

Double-stranded RNA was transcribed *in vitro* from DNA templates as described previously^25^. In brief, two partially complementary DNA oligomers with the following sequence were used for dsRNA synthesis: dsRNA_A, 5’-gggagaatgtcgaatgggtattccacagacgagaatttccgctatctcatctcgtgcttcagggccagggtgaaaatgtacatccaggtggagcctgtgctggactacctgacctttctgcctgcagaggtgaaggagcagattcagaggacagtctctccc-3’; dsRNA_B, 5’-gggagaatgaacacgattaacatcgctaagaacgacttctctgacatcgaactggctgctatcccgttcaacactctggctgaccattacggtgagcgtttagctctctccc. The lyophilized DNA oligomers were synthesized by Integrated DNA Technologies (Coralville, IA, USA) and resuspended in dH_2_O. RNA was transcribed *in vitro* by combining 200 ng DNA template with T7 RNA Polymerase (NEB #M0251S), 1X NTPs (N0450S), 5 mM DTT, and 20 U RNase Inhibitor (NEB #M0314S); the transcription reaction was incubated for 18 h at 37 °C. The RNA was isolated from the transcription reaction by following the manufacturer’s instructions for the RNA Clean & Concentrator Kit (Zymo #R1013). The RNA purity was assessed using the Agilent RNA ScreenTape (Agilent #5067-5576 and Agilent #5067-5577). Purified ssRNA was combined at a 1.44:1 molar ratio (dsRNA_A:dsRNA_B) and hybridized by heating to 85 °C for 2 min before cooling to 25 °C. Exogenous dsRNA (100 ng dsRNA/cm^2^ plate surface area) was transfected into HEK293T cells using the JetPEI transfection system (see above).

### Mycoplasma Testing

All cell lines were routinely tested for mycoplasma every month using the MycoAlert Mycoplasma Detection Kit (Lonza #LT07-318) following the manufacturer’s protocol. New cell lines were quarantined and tested for mycoplasma twice at least one week apart before moving out of quarantine. DAPI staining was sporadically used to further confirm that mycoplasma contamination was not present in cultured cells.

### Human Brain Models

Control human brain tissue samples were obtained from the UCSD ALS repository, which is compliant with HIPAA informed consent procedures and approved by the Institutional Review Board (CA IRB #120056). The brain tissue was dissected in an autopsy suite (usually with a short postmortem interval of less than 6 h) and either immediately frozen for biochemical studies or fixed in neutral buffered formalin for 2 weeks. Frontal cortex derived from three mid-age and three old-age patient samples (Table S8) were used for this study. For dsRNA-seq and PKR CLIP experiments, only cases 26, 131, 65, and 103 were used because the other two samples failed sequencing quality control. Human brain tissue is available via the Target ALS Multicenter Human Postmortem Tissue Core and can be requested at the following webpage: https://www.targetals.org/resource/human-postmortem-tissue-core/.

### Immunofluorescence

Ibidi μ-slide 18-well chambers (Ibidi #81811) were coated as described above with PDL/PLO/laminin or 1X Matrigel for transdifferentiated and iPSC-derived neurons, respectively. Cells were dissociated and transferred to the Ibidi μ-slides at least 48 h prior to immunofluorescence. The samples were fixed with 4% (v/v) paraformaldehyde (Fisher #50980492) at 4 °C with gentle rocking for 45 min. The fixed cells were thoroughly washed three times for 5 min each with 1X dPBS. The samples were then permeabilized with 1X dPBS with 0.1% (v/v) Triton X-100 (Sigma #X100-500ML) and 5% (v/v) donkey serum (Jackson ImmunoResearch #017-000-121) for 45 min at 25 °C. The samples were washed three times for 1 min each with 1X dPBS with 0.1% (v/v) Triton X-100 (dPBST). Primary antibodies were added to antibody solution (1X dPBST with 0.1% (w/v) bovine serum albumin, Fisher #10773877) at the concentrations listed in Table S9 and incubated for 24 h with gentle rocking at 4 °C. The cells were then washed an additional three times with dPBST for 5 min each. Secondary antibodies were diluted in antibody solution (see Table S9) and incubated with the samples for 1 h with gentle rocking at 25 °C. Following this incubation, the cells were washed for 5 min three times with 1X dPBS, and Prolong Diamond Antifade Mountant with DAPI (Thermo #P36971) was used for long-term storage of the slides at 4 °C. The immunofluorescence samples were imaged on a Zeiss LSM-880 microscope with 405, 488, 561, and 633 nm lasers using a 63X/1.4 DIC oil immersion objective. Confocal imaging data were acquired as single slices and line averaging was used to improve signal-to-noise ratio. For Airyscan imaging data, processing of all channels was performed in Zen software using the default processing settings.

Image quantification and processing was performed in ImageJ. All images were taken with the same acquisition settings and processed with the same manipulations for each panel unless otherwise indicated. For intensity profiles, the “Plot Profile” function was used on a line drawn through the cytoplasm and nucleus of the indicated cell. For stress granule counting, samples were anonymized for quantification, and stress granules were manually counted in each condition.

### Immunofluorescence and Fluorescence In Situ Hybridization (IF-FISH)

FISH probes were synthesized by IDT with the following sequences and fluorophore modifications^76^: MT-ND6, 5’-/Cy3/ccatgcctcaggatactcctcaatagccatcgctgtagtatatccaaaga-3’; and MT-RNR1, 5’-/Cy3/tggctggcacgaaattgaccaaccctggggttagtatagcttagttaaac-3’. IF-FISH experiments were performed as described above for IF through the final washing steps before mounting. After three washes, neurons were fixed again in 4% (v/v) paraformaldehyde for 10 min at 25 °C with gentle rocking. Meanwhile, a homemade humidity chamber was pre-warmed to 37 °C. The re-fixed cells were washed with dPBST twice for 5 min each and FISH Wash Buffer (2X SSC with 10% formamide) for 5 min at 25 °C with gentle rocking. Following the washes, 1 μg of fluorescent FISH probe in FISH Hybridization Buffer (2X SSC, 10% dextran sulfate, 0.2% (w/v) BSA) was added to each sample, and the hybridization was performed for 24 h at 37 °C in the pre-warmed humidity chamber. The hybridized samples were then washed with FISH Wash Buffer twice for 5 min each at 37 °C, and an additional two washes were performed with 2X SSC at 37 °C for 5 min each. Finally, the samples were mounted with Prolong Diamond Antifade Mountant with DAPI and imaged as described above.

### Mitochondria Live-Cell Tracking

Glass coverslips were pretreated with 10% (v/v) nitric acid overnight at room temperature and washed 10 times over a period of 3 days, followed by a final rinse in 70% ethanol to increase neuronal adherence. Following pretreatment, the coverslips were coated with PDL/PLO/Laminin as described above. Neurons were passaged from differentiation plates onto the coverslips, and the area surrounding each well was filled with sterile dPBS to maintain humidity. Neurons were incubated on the coverslips for 48 h at 37 °C prior to imaging. For staining, the neurons were treated with 160 nM TMRM (Thermo #I34361) and 60 nM MitoTracker Green (Thermo #M7514) for 20 min at 37 °C. Live-cell imaging was performed on a Zeiss LSM780 laser scanning confocal microscope outfitted with a heated stage and laminar-flow perfusion system; videos were acquired with a 40x objective. The cells were continuously perfused with Hibernate E media (Thermo #A1247601) supplemented with TMRM; frames were acquired every 40 s using the Definite Focus setting. Antimycin A (Sigma-Aldrich #A8674) and rotenone (Sigma-Aldrich #R8875) (5 μM each) were added ∼5 minutes after the start of the video, and acquisition continued for another 5 minutes after Antimycin A and rotenone perfusion. Videos were analyzed using ImageJ to track MitoTracker Green and TMRM intensity in individual mitochondria regions-of-interest; the intensity values were normalized to the first time point and ψ_m_ was determined by dividing the TMRM intensity by the MitoTracker Green intensity.

### RNA Isolation

Flash-frozen cell pellets were thawed on ice and resuspended in 1 mL Trizol (Thermo #15596026). After reacting with TRIzol for 5 min, 200 μL chloroform (Fisher #C298-500) was added to the samples, which were then vigorously vortexed for 15 s. The TRIzol and chloroform mixture was centrifuged at 21000 x g for 15 min at 4 °C. The upper aqueous phase was carefully extracted from the centrifuged samples, and the aqueous phase was combined with an equal volume of 70% (v/v) ethanol. This mixture was then loaded onto the RNeasy purification spin column (Qiagen #74104), and the manufacturer’s instructions were followed for the remainder of the RNA isolation protocol. For brain tissue, the same protocol was used except flash-frozen brain was resuspended in TRIzol LS (ThermoFisher #10296010).

### dsRNA Dot Blots

Purified total RNA was quantified using a ThermoScientific NanoDrop Eight; the RNA was diluted in increasing concentrations for loading onto the dot blot. Diluted RNA was manually loaded onto an Amersham Hybond-N^+^ nylon membrane (Cytiva #RPN203B), air-dried, and UV crosslinked using a UVP CL-1000 Ultraviolet Crosslinker (4000 μJ exposure). The cross-linked membrane was blocked in 5% (w/v) milk powder (Apex Bioresearch #20-241) in 1X TBST (1X TBS, Fisher #AAJ62662K7; 0.1% (v/v) Tween-20, Sigma #P9416) for 30 min at 25 °C with gentle rocking. After blocking, the membrane was then incubated with the J2 antibody (Scicons #10010200, diluted 1:1000) for 18 h at 4 °C with gentle rocking. Following three washes with 1X TBST, the membrane was incubated with an HRP-conjugated secondary antibody (Cell Signaling Technology #7076, diluted 1:1000) for 2 h at 25 °C with gentle rocking. After an additional three washes with 1X TBST, the HRP-conjugated secondary antibodies were developed with ECL Western Blotting Substrate (Life #32106) and imaged on an Azure 600 Western Blot Imager (Azure Biosystems). Total RNA was visualized using Methylene Blue (Sigma #M9140-25G). Dot blots were analyzed using the ImageJ Gel Analysis function.

### RNA-seq

Total RNA was isolated as described above and prepared for RNA sequencing using the Illumina Stranded mRNA Prep kit (Illumina #20040534) and the IDT for Illumina DNA/RNA UD Indexes (Illumina #20040553) as described in the manufacturer’s protocol. The sequencing libraries were analyzed using a D1000 ScreenTape (Agilent #5067-5582) with an Agilent 4150 TapeStation. Pooled libraries were sequenced using either an Illumina NextSeq 2000 or NovaSeq 6000 with PE50 settings. Sequencing reads were processed and analyzed using the NF-Core pipeline^77^.

### dsRNA-seq

At least 5 x 10^6^ cells were pelleted for J2 (dsRNA) immunoprecipitation and sequencing. Flash-frozen cells were resuspended in 1 mL TRIzol (Thermo #15596026) and incubated at 25 °C for 5 min. Chloroform (200 μL) was added and the samples were vigorously vortexed for 15 s. The samples were then centrifuged at 18000 x g for 15 min at 4 °C, and the upper aqueous phase containing total RNA was carefully extracted to a new tube. Isopropanol (500 μL) and glycoblue (1 μL; Thermo #AM9515) were added to the aqueous phase. The RNA was precipitated by incubating at -80 °C for 10 min, and the samples were centrifuged at 12000 x g for 10 min at 4 °C to pellet the RNA. After removing the supernatant fraction, the RNA pellet was resuspended in 1 mL 75% (v/v) ethanol and precipitated again. The washed pellet was air-dried for 10 min, resuspended in 50 μL dH_2_O, and analyzed using an RNA ScreenTape (Agilent #5067-5576) on an Agilent TapeStation 4150 and ThermoFisher Nanodrop to determine the RNA concentration.

Total RNA (50 μg) was digested with 2 U DNase (ThermoFisher #AM2238) in the presence of RNase Inhibitor (NEB #M0314S) for 20 min at 37 °C. Meanwhile, J2-conjugated beads were prepared by washing M-280 Sheep Anti-Mouse IgG Dynabeads (ThermoFisher #11201D) with J2-B Buffer (20 mM Tris-HCl, pH 7.5; 150 mM NaCl; 0.2 mM EDTA, pH 8.0; 0.2% (v/v) Tween-20) prior to binding with 5 μg Scicons J2 Antibody (Scicons #10010200). The J2-conjugated beads were reacted with the DNase-digested RNA for 18 h at 4 °C with rotation; the “input” sample was removed just before this step for total RNA sequencing.

Following J2 immunoprecipitation, the beads were washed three times in J2-B Buffer and resuspended in 150 μL. RNA was phase-separated from the beads by adding 450 μL TRIzol LS (ThermoFisher #10296028), incubating for 5 min, then adding 120 μL chloroform with 15 s vigorous vortexing to mix. The samples were centrifuged at 18000 x g for 15 min at 4 °C, and the upper aqueous layer containing the RNA was extracted to a new tube. The immunoprecipitated RNA was further purified using the Qiagen RNeasy Mini Kit (Qiagen #74104) by adding 375 μL 70% (v/v) ethanol, vortexing for 5 s, and loading onto the RNeasy column. The manufacturer’s directions were followed for the remainder of the RNeasy column purification, and the eluted RNA was quantified with the Agilent TapeStation.

RNA libraries were prepared following the Illumina mRNA Stranded Prep (Illuina #20040534) workflow with a few modifications: (1) the poly(A) selection step was bypassed; and (2) prior to fragmentation, 1 μg RNA was combined with 2X EPH3 and dH_2_O to a final volume of 17 μL and concentration of 1X EPH3 solution. The IDT for Illumina UD Indexes (Illumina #20027213) were used for library amplification, and the samples were sequenced using a NextSeq 2000 (Illumina, San Diego, CA) following the manufacturer’s protocol. The reads were demultiplexed using the Illumina BaseSpace application DRAGEN BCL Convert v4.2.7, and the RNA-seq reads were further processed with the NF-Core pipeline^77^.

### eCLIP-seq

CLIP experiments were performed essentially as described before^34,35^. In brief, at least 1 x 10^7^ neurons were UV-crosslinked, lysed, and reacted with antibody-conjugated Dynabeads. The following antibodies were used for eCLIP: 1 μg Rabbit α-PKR (Cell Signaling Technology #12297S); 10 μg Rabbit α-PNPT1 (Bethyl #A303-917A); and 2 μg Rabbit α-SUV3 (Bethyl #A303-056A). Immunoprecipitated RNA was repaired and size-selected using gel electrophoresis and transferred to a nitrocellulose membrane. Libraries were prepared from the size-selected RNA and sequenced using a NextSeq 2000 instrument. Analysis of eCLIP peaks was performed using the Skipper pipeline^78^ through the web portal at https://rbp-ark.com/.

### Riboseq

Riboseq was performed essentially as described previously^79^. In brief, cells were harvested and lysed in 100 μg/mL cycloheximide to stall translation and arrest mRNAs on ribosomal footprints. Ribosomal-shielded RNA was precipitated by sucrose gradient centrifugation. The resulting fragments were gel-purified, repaired for library preparation, and converted into cDNA. The library was amplified and sequenced using a NextSeq 2000. The Riboseq reads were analyzed using the xtail package as described previously^80,81^.

### Western Blotting

At least 1 x 10^5^ cells were pelleted and stored at -80 °C for Western blotting. Cell pellets were resuspended in 200 μL RIPA Lysis and Extraction Buffer (Thermo #89900) supplemented with 2X Halt Protease and Phosphatase Inhibitor Cocktail (#Thermo #78440). The resuspended cells were incubated with rotation at 4 °C for 15 min, and they were further lysed by sonicating for 10 min (30 s/30 s off cycles). Lysed cells were centrifuged at 16000 x g for 15 min at 4 °C. The supernatant fraction was transferred to a new tube and combined with 4X NuPAGE LDS Sample Buffer (Thermo #NP0007). The samples were then heated to 70 °C for 10 min and loaded onto 15-well NuPAGE 4-12% Bis-Tris Mini Protein Gels (Life #NP0323BOX). Gels were electrophoresed in the Novex XCell SureLock Mini-Cell Electrophoresis System (Thermo #EI0001) at 150 V for 75 min. The samples were then transferred to Immobilon-FL PVDF membranes (Sigma #IPFL00010) pre-soaked in methanol. The electrophoresis was performed in Transfer Buffer (1X NuPAGE Transfer Buffer, Life #NP0006; 10% (v/v) methanol, Fisher #A411-4) for 16 h at 40 V. Once the samples were transferred to the PVDF membrane, the membranes were blocked in 5% (w/v) BSA in 1X TBST (1X TBS, Fisher #AAJ62662K7; 0.1% (v/v) Tween-20, Sigma #P9416) for 30 min at 25 °C. The following primary antibodies were diluted in 5% (w/v) milk powder dissolved in 1X TBST: 1:1000 Rabbit Anti-Phospho-eIF2α (Abcam #ab32157); and 1:1000 Rabbit Anti-eIF2α (Cell Signaling Technology #9722S). Once diluted, the antibodies and blocked membrane were incubated overnight at 4 °C with gentle rocking. The membrane was then washed thoroughly three times with 1X TBST for 5 min each wash at 25 °C. The IRDye® 680RD Goat anti-Rabbit IgG Secondary Antibody was diluted 1:5000 in 5% (w/v) BSA in 1X TBST and incubated with the membrane for 1 h at 25 °C with gentle rocking. After an additional three washes with 1X TBST, the fluorescent antibodies were imaged on an Azure 600 Western Blot Imager (Azure Biosystems). The Western blot images were analyzed using ImageJ to determine the intensity of individual bands, which were normalized to the loading control. The relative expression was plotted using GraphPad Prism 10, and Welch’s t-test was used to calculate significance in Excel.

### J2 Immunoprecipitation and Mass Spectrometry

At least 5 x 10^6^ cells were pelleted for J2 (dsRNA) immunoprecipitation and mass spectrometry. Frozen cell pellets were resuspended in IP Lysis Buffer (50 mM Tris-HCl, pH 7.4; 150 mM NaCl; 1 mM EDTA, pH 8.0; 0.5% (v/v) IGEPAL-CA630) supplemented with fresh protease inhibitor cocktail. The cells were then sonicated for 5 min (30 s on/30 s off), and the lysate was centrifuged at 14000 x *g* for 10 min at 4° C. Following centrifugation, the supernatant fraction was transferred to a new tube, and the protein content of each sample was estimated using the Pierce BCA Protein Assay Kit (Thermo Scientific #23225). Approximately 5 mg protein was used for immunoprecipitation. Five micrograms per sample of J2 (dsRNA) antibody (Scicons #10010200) or Mouse Anti-IgG (Invitrogen #026502) were incubated with washed Sheep Anti-Mouse M-280 Dynabeads (Invitrogen #11201D) for 45 minutes at 25 °C with end-over-end rotation. The antibody-conjugated beads were washed with IP Lysis Buffer three times; the washed beads were combined with the lysed protein and incubated for 16 h at 4 °C with rotation. The beads were then washed three times with IP Lysis Buffer and three times with IP Wash Buffer (50 mM Tris-HCl, pH 7.4; 150 mM NaCl); any residual liquid was removed, and the beads were frozen at -20 °C prior to mass spectrometry preparation.

Proteins bound to the antibody-conjugated beads were reduced with 10 mM TCEP (Sigma #4706) for 30 min, alkylated with 15 mM freshly prepared iodoacetamide (IAA; Sigma #I6125) for 45 min in the dark, and then treated with 10 mM DTT for 15 min in the dark. The protein was then purified via methanol-chloroform precipitation and solubilized in 100 mM TEAB (Sigma #T7408). For digestion, 200 ng LysC endoprotease (NEB #P8109S) and 400 ng trypsin (NEB #P8101S) were added to the samples, which were then incubated at 37 °C with shaking at 300 rpm for 16 h. The digestion was terminated by adding formic acid (Sigma #F0507) to a final concentration of 1%.

The digested peptides were desalted using the Stage-Tip method and then analyzed by an LC-MS/MS setup. The setup included a timsTOF Pro 2 mass spectrometer (Bruker Daltonics, Bremen, Germany) coupled to a nanoElute 2 nano-LC system (Bruker Daltonics, Bremen, Germany), using a CaptiveSpray ion source. The peptides were eluted through a reversed-phase C18 column (PepSep, 25 cm x 150 µm, 1.5 µm) maintained at 50 °C over a 50 min gradient of 5-35% solvent B (acetonitrile in 0.1% formic acid) at a flow rate of 500 nL/min. The peptides were analyzed using data independent acquisition (dia) Parallel Accumulation-Serial Fragmentation (PASEF) mode. The isolation windows for diaPASEF were determined based on data dependent acquisition (dda)-PASEF data. The settings for ddaPASEF included the following: a ramp time and an accumulation time of 75 ms each, one MS scan, and 10 PASEF MS2 ramps per acquisition cycle. The MS survey scan covered a mass-to-charge (m/z) ratio range of 100-1700 m/z and ion mobility (1/k0) range of 0.6-1.6 V s/cm^2^. Precursors with up to 5 charges were selected, with an active exclusion time of 0.4 min. The quadrupole isolation width was set at 2 m/z for m/z values below 700 and 3 m/z for m/z values above 800, with linear interpolation for m/z values of 700-800. For diaPASEF, the isolation windows were designed to encompass the precursor distribution across the m/z-1/k0 plane as defined by the ddaPASEF data, ranging from 323.6 to 1221.6 m/z and 0.7 to 1.34 V s/cm^2^ in 1/k0. Each 75 ms diaPASEF scan spanned a 40 Da mass width with a 1 Da mass overlap, and the number of windows varied to cover the entire ion mobility spectrum (creating 23 windows for 10 diaPASEF scan cycles).

The diaPASEF raw files were processed using a library-free search approach in DIA-NN version 1.8.1. An *in silico* digestion of a UniProt reviewed Homo sapiens database (December 2022) was employed to predict the spectral library. The search parameters applied included the following: tryptic digestion with up to two missed cleavages, carbomidomethylation of cysteine as static modification, oxidation of methionine, and N-terminal acetylation as variable modifications. The precursor false discovery rate (FDR) threshold was set at 1%. The quantification strategy for precursors was set to Robust LC (high precision) and cross-run normalization was set to RT-dependent. Match between run enabled for the protein group matrix file generated by DIA-NN, filtered at 1% FDR using global q-values for protein groups and global and run-specific q-values for precursors, was used for downstream and statistical analysis. The raw intensities for each identified protein group were used to calculate and visualize fold changes and p-values in Excel. dbSTRING networks were constructed in CytoScape with an interaction confidence cutoff of 0.95. Top gene ontology terms were calculated using ShinyGO v0.82^82^, and these terms were filtered to remove any gene ontology groups with <60 genes.

### G3BP1 Immunoprecipitation and Mass Spectrometry

At least 5 x 10^6^ cells were treated with PKR inhibitor or DMSO as described above and pelleted for G3BP1 immunoprecipitation and mass spectrometry. Frozen cell pellets were resuspended in IP Lysis Buffer (50 mM Tris-HCl, pH 7.4; 150 mM NaCl; 1 mM EDTA, pH 8.0; 0.5% (v/v) IGEPAL-CA630) supplemented with fresh protease inhibitor cocktail. The cells were then sonicated for 5 min (30 s on/30 s off), and the lysate was centrifuged at 14000 x *g* for 10 min at 4° C. Following centrifugation, the supernatant fraction was transferred to a new tube, and the protein content of each sample was estimated using the Pierce BCA Protein Assay Kit (Thermo Scientific #23225). Approximately 10 mg protein was used for immunoprecipitation. Five micrograms per sample of Rabbit anti-G3BP1 antibody (MBLI #RN048PW) were incubated with washed Sheep Anti-Rabbit M-280 Dynabeads (Invitrogen #11203D) for 45 minutes at 25 °C with end-over-end rotation. The antibody-conjugated beads were washed with IP Lysis Buffer three times; the washed beads were combined with the lysed protein and incubated for 16 h at 4 °C with rotation. The beads were then washed three times with IP Lysis Buffer and three times with IP Wash Buffer (50 mM Tris-HCl, pH 7.4; 150 mM NaCl); any residual liquid was removed, and the beads were frozen at -20 °C prior to mass spectrometry preparation. G3BP1 AP-MS samples were submitted to the Sanford Burnham Prebys Proteomics Core for tryptic digestion, desalting, mass spectrometry, and data processing.

### Mitochondrial Stress Test

Seahorse XFe96/XF Pro Cell Culture Microplates (Agilent #103794-100) were coated with PDL/PLO/Laminin as described above. Following the coating, neurons were detached from their differentiation plate and counted with a Countess 3 Automated Cell Counter (Thermo Fisher Scientific); 4.5 x 104 neurons were seeded per well and incubated for 48 h at 37 °C prior to the assay. For respirometry measurements, the neuron’s medium was exchanged with XF DMEM Base Medium (pH 7.4) with no Phenol Red (Agilent 103575-100) supplemented with 5 mM D-glucose (Fisher #D16-500) and 1 mM sodium pyruvate (Life Technologies 11360070). The oxygen consumption rate was measured using an Agilent Seahorse XFe96 Analyzer (Agilent) after injections of the following compounds at concentrations varying from 0.5-5 μM: oligomycin (Sigma-Aldrich #75351), carbonyl cyanide 4-(trifluoromethoxy) phenylhydrazone (FCCP) (Sigma-Aldrich #C2920), and rotenone with antimycin A. NucBlue dye (Thermo Fisher Scientific) was included to stain nuclei for cell counting after the assay. The cell count was normalized using the custom written FluxNormalyzer macro, and the respirometry values were analyzed as described previously^83,84^.

### Statistics & Reproducibility

No statistical method was used to predetermine sample sizes. Most experiments were not anonymized or randomized during experiments; however, sample identity was blinded for stress granule quantification. Outlier data points were only excluded if they tested as a significant outlier using Grubb’s test. For simple statistical comparisons, a two-tailed Welch’s t-test (i.e. a t-test assuming a normal distribution but with unequal variances) was performed in most cases, and the degrees of freedom for significance scoring was approximated using Satterthwaite’s correction. We did not confirm normal distributions for each data distribution. Certain data analysis pipelines (e.g. ShinyGO) have their own statistical calculations built into the software, and the original work will provide greater detail about the statistical power of their calculations. For imaging data and blots, we performed each experiment at least in triplicate; certain imaging experiments (e.g. Map2/G3BP1/J2 staining) were performed over ten times, and we observed strong reproducibility across many different biological and technical replicates. The images shown in the manuscript are a representative sample of each experiment.

## Supplemental Figures

**Figure S1:**
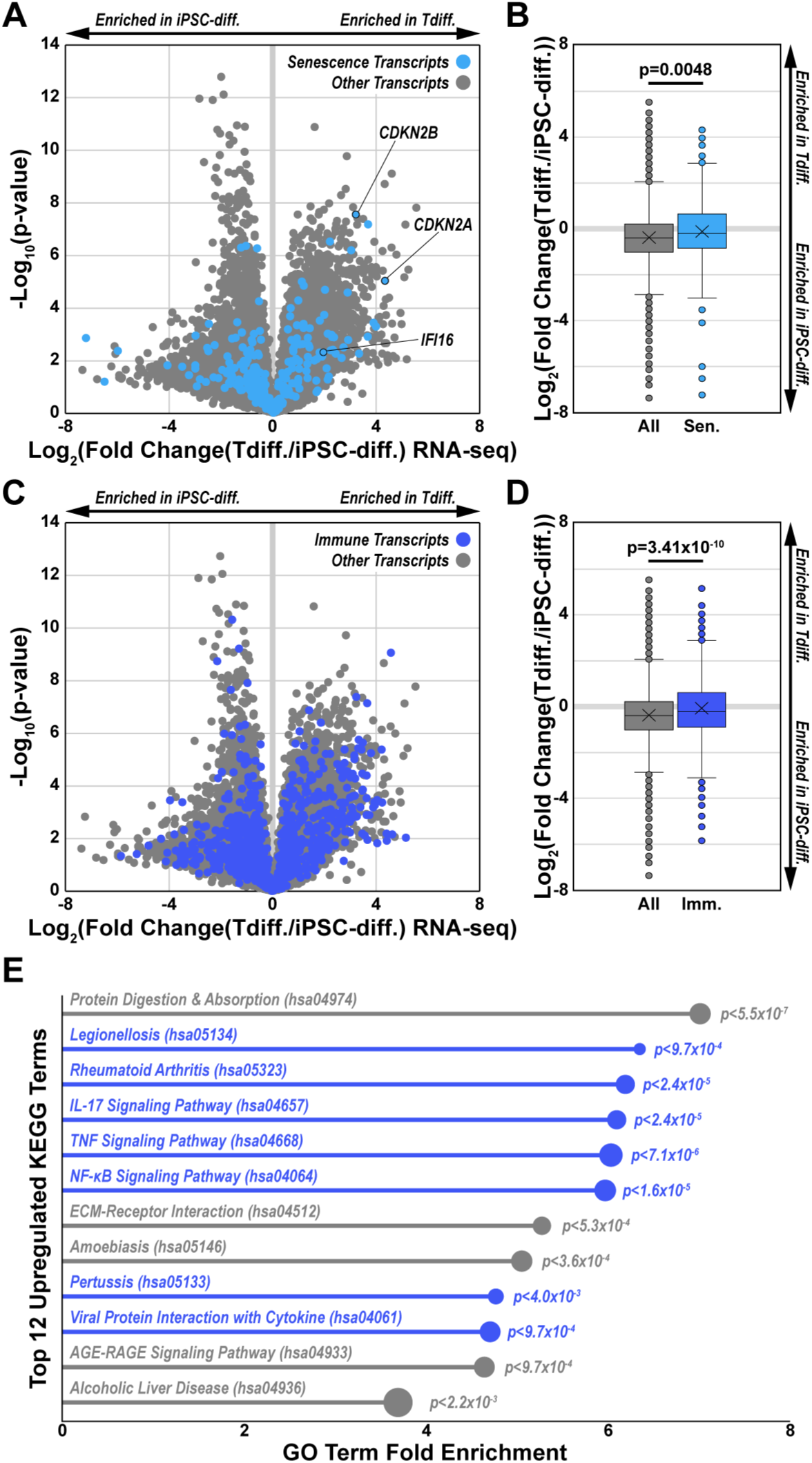
Aging- and immune-linked pathways are upregulated in transdifferentiated neurons. **(A)** Volcano plot of RNA-seq in transdifferentiated neurons (Tdiff.1/2/4/5) versus iPSC-derived neurons (n=3 replicates per line; N=4 lines per cohort). Light blue data points denote senescence-associated transcripts; all other transcripts are colored gray. Key senescence transcripts are highlighted on the plot. **(B)** Boxplot of the Log_2_(Fold Change) of all transcripts (gray) and senescence-linked transcripts (light blue) in transdifferentiated neurons versus iPSC-derived neurons (n=3 replicates per line; N=4 lines per cohort). From top to bottom, the box plot lines denote the 75^th^, 50^th^, and 25^th^ percentile of data values; error bars denote the range of non-outlier values, the “X” denotes the mean, and any indicated points are outliers. Statistics were calculated using a two-tailed Welch’s t-test for each treatment condition compared to the control. **(C)** Same as (A) but with the gene ontology pathway “immune system process” (GO:0002376) highlighted in dark blue. **(D)** Same as (B) but with the “immune system process” pathway. **(E)** Lollipop plot of the top 12 most enriched KEGG pathways that were upregulated >4-fold in transdifferentiated neurons compared to iPSC-derived neurons as calculated by ShinyGO v0.82. The size of the lollipop circle corresponds to the number of genes in the pathway, and the p-value is the enrichment false discovery rate. Pathways that are linked to the immune response are highlighted dark blue.

**Figure S2:**
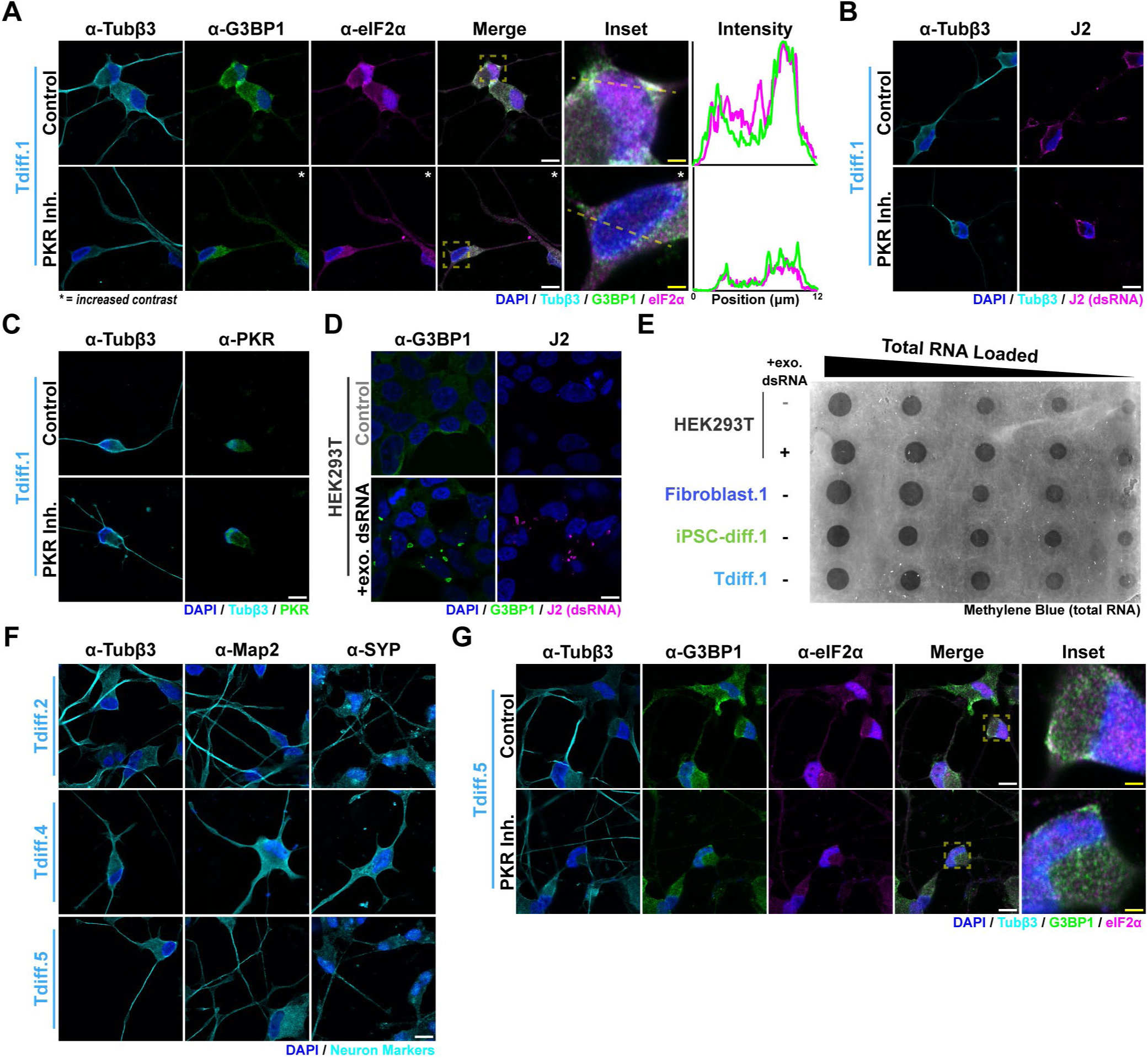
Stress granules are resolved by PKR inhibition in transdifferentiated neurons. **(A)** Confocal immunofluorescence images of Tubβ3 (cyan), G3BP1 (green), eIF2α (magenta), and DAPI (blue) in Tdiff.1 neurons with and without PKR inhibitor treatment. The contrast of the indicated panels was increased for better visualization in the figure. The yellow dashed box indicates the inset region, and the yellow dashed line indicates the plot profile in the intensity panels. White Scale Bar = 10 μm, Yellow Scale Bar = 2 μm. **(B)** Confocal immunofluorescence images of Tubβ3 (cyan), J2 (magenta), and DAPI (blue) in Tdiff.1 neurons with and without PKR inhibitor treatment. **(C)** Same as (B) but with PKR (green) instead of J2 staining. **(D)** Confocal immunofluorescence images of G3BP1 (green), J2 (magenta), and DAPI (blue) in HEK293T cells treated with and without exogenous dsRNA. **(E)** Methylene blue (total RNA) staining of the dot blot in Figure 1F. **(F)** Confocal immunofluorescence images of neuronal markers (cyan) and DAPI (blue) in different transdifferentiated neuron lines (Tdiff.2/3/5). Scale Bar = 10 μm. **(G)** Same as (A) but for Tdiff.5.

**Figure S3:**
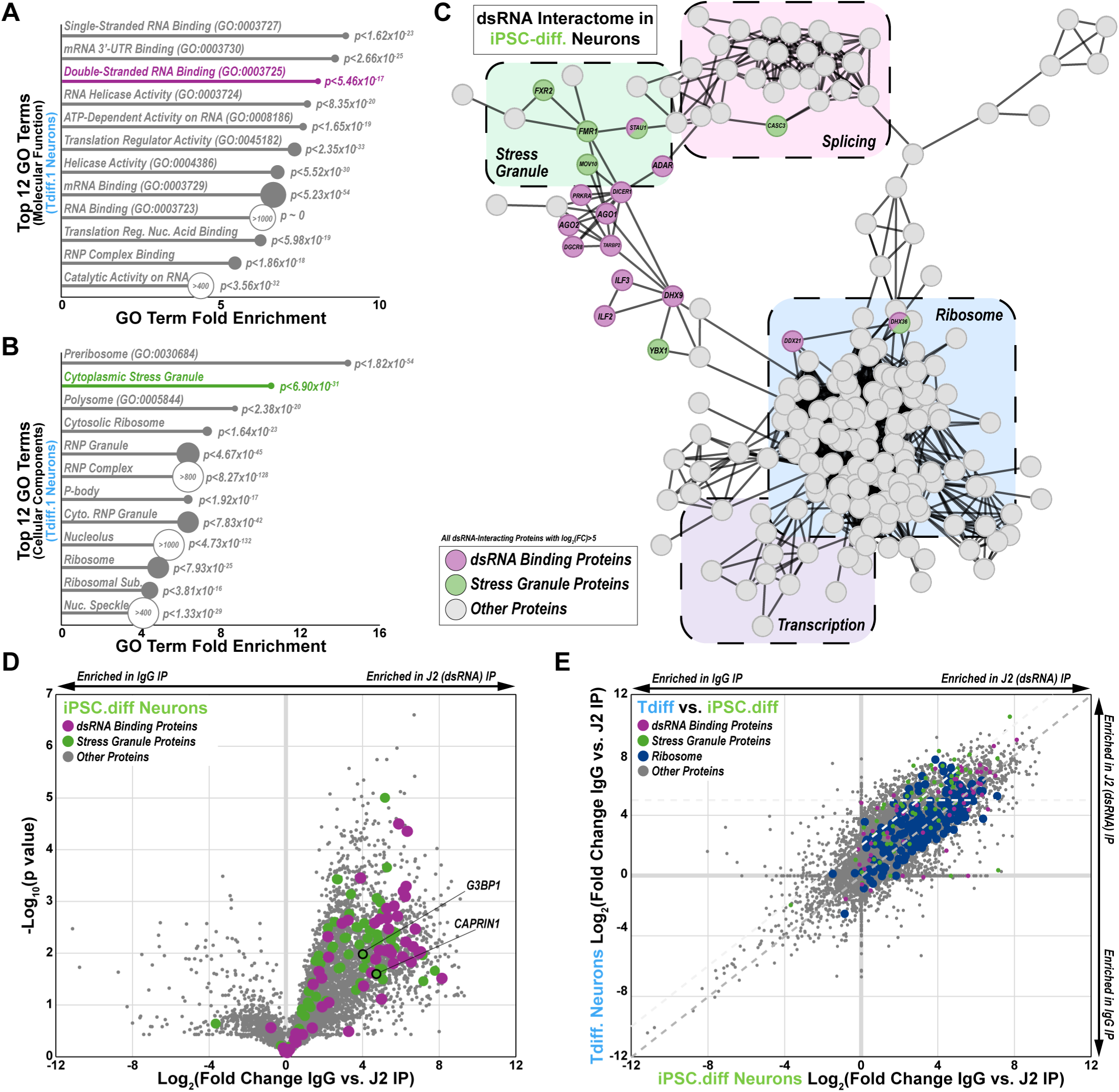
Ribosomal proteins are not over-enriched in the dsRNA interactome of transdifferentiated neurons. **(A)** Lollipop plot of the top 12 Molecular Function Gene Ontology terms for the Tdiff.1 dsRNA interactome; only proteins with a log_2_(FC)>5 for the J2/IgG pulldowns were included. The circle size denotes the number of genes in each term list, except as indicated. **(B)** Same as A but for the top 12 Cellular Components Gene Ontology terms. **(C)** dbSTRING network of proteins that were immunoprecipitated by dsRNA (J2) in iPSC-derived neurons (iPSC-diff.1). Only significantly interacting proteins (log_2_(FC)>5 for J2/IgG pulldown) that clustered within the main network are shown. The superposition of distinct groups of proteins (e.g. ribosome, transcription, etc.) is approximate. Nodes of dsRBPs (purple; GO:0003725) and stress granule proteins (green; GO:0010494) are labeled on the network. **(D)** Volcano plot of the log_2_(FC) of dsRNA-interacting proteins in iPSC-derived neurons (iPSC-diff.1). Proteins of interest are labeled as in (C), and the canonical stress granule markers G3BP1 and Caprin1 are indicated. **(E)** Fold change versus fold change plot of J2/IgG pulldown in transdifferentiated neurons and isogenic iPSC-derived neurons. dsRBPs and stress granule proteins are highlighted as in (C), and ribosomal proteins (GO:0005761) are also indicated in orange. The gray dashed line denotes the y=x line whereas the light gray dashed lines denote log_2_(FC)>5 for the Tdiff.1 pulldown and log_2_(FC-Tdiff.1) −log_2_(FC-iPSC.diff.1)>2.

**Figure S4:**
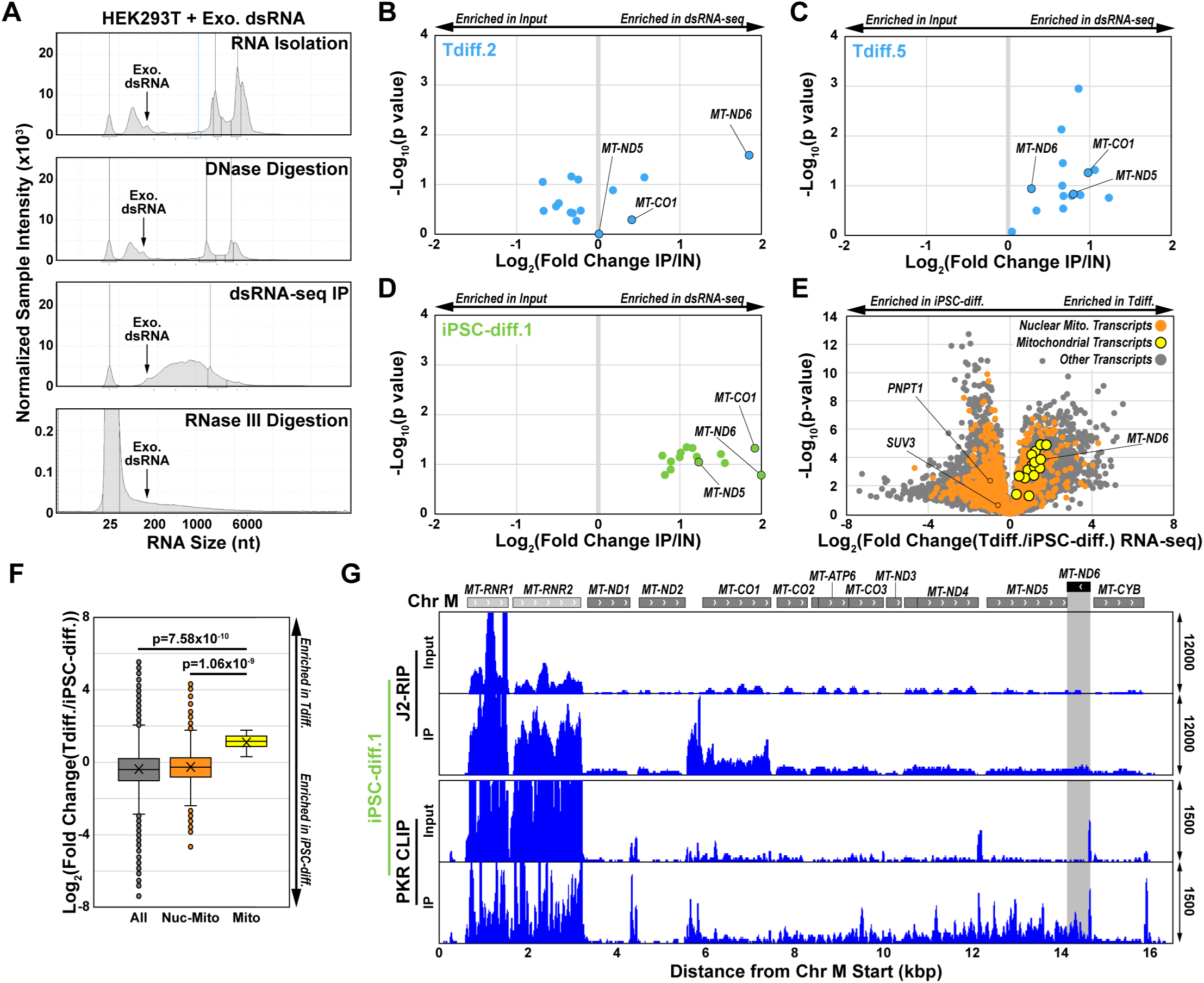
PKR does not strongly bind to MT-dsRNA in iPSC-derived neurons. **(A)** Screen captures from the Agilent TapeStation Analysis Software v4.1.1 of RNA ScreenTapes loaded with samples from throughout the dsRNA purification protocol in HEK293T cells. Exogenous (exo.) dsRNA of a known size (∼150 bp) was tracked throughout the purification. **(B)** Volcano plot of the log_2_(Fold Change) of all MT-RNA transcripts identified by dsRNA-Seq in the input and immunoprecipitation (IP) samples of the Tdiff.2 transdifferentiated neuron line. **(C)** Same as (B) but for Tdiff.5 transdifferentiated neurons. **(D)** Same as (B) but for iPSC-diff.1 neurons. **(E)** Volcano plot of RNA-seq in transdifferentiated neurons (Tdiff.1/2/4/5) versus iPSC-derived neurons (n=3 replicates per line; N=4 lines per cohort). Orange data points denote nuclear-encoded mitochondrial genes whereas yellow data points denote MT-RNA transcripts. The MT-ND6, SUV3, and PNPT1 transcripts are indicated on the plot. **(F)** Boxplot of the Log_2_(Fold Change) of all transcripts (gray), nuclear-encoded mitochondrial transcripts (orange), and MT-RNA transcripts (yellow) in transdifferentiated neurons versus iPSC-derived neurons (n=3 replicates per line; N=4 lines per cohort). **(G)** Visualization of representative mitochondrial sequencing reads detected by dsRNA-Seq and PKR CLIP in the Tdiff.1 transdifferentiated neuron line. The shaded region denotes the reverse-transcribed *MT-ND6* gene. The range of the y-axis values is denoted on the right side of the genomic track and was fixed for each pair of input and IP samples.

**Figure S5:**
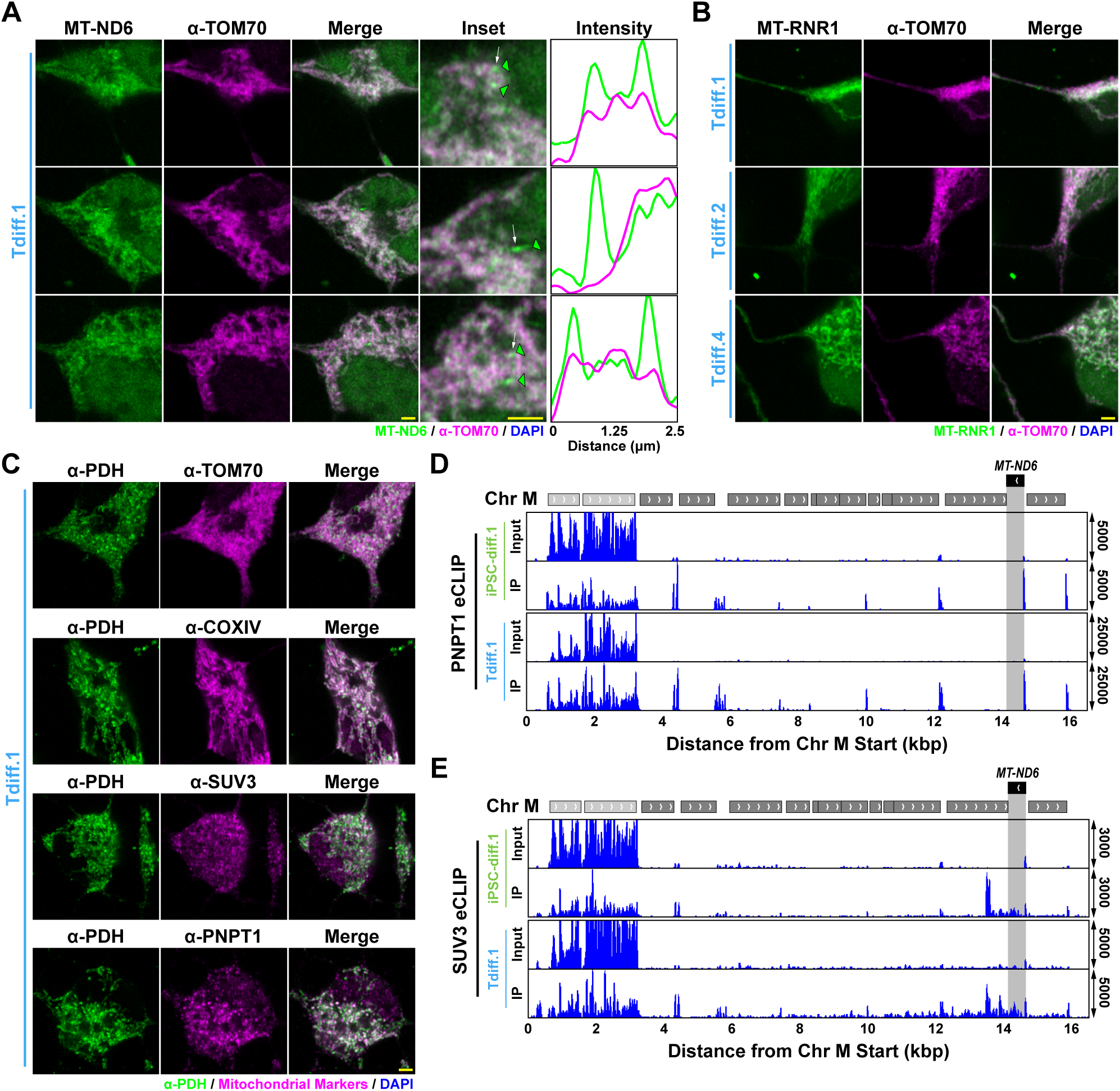
Mitochondrial degradasome proteins bind to MT-dsRNA in transdifferentiated neurons. **(A)** Additional airyscan immunofluorescence-FISH images of the mitochondrial marker TOM70 (magenta) and MT-ND6 transcript (green) in Tdiff.1 neurons. Green arrowheads denote cytoplasmic MT-ND6 puncta; the white arrows denote the intensity profiles in the adjacent plot. Yellow Scale Bar = 2 μm. **(B)** Same as (A) but for the MT-RNR1 transcript in various transdifferentiated neuron lines (Tdiff.1/2/4). **(C)** Airyscan immunofluorescence images of the indicated mitochondrial proteins in Tdiff.1 neurons. Yellow Scale Bar = 2 μm. **(D)** Visualization of representative mitochondrial sequencing reads detected by PNPT1 eCLIP (n=2 replicates) in the indicated neuronal lines. The shaded region denotes the reverse-transcribed *MT-ND6* gene. **(E)** Same as (D) but for SUV3 eCLIP.

**Figure S6:**
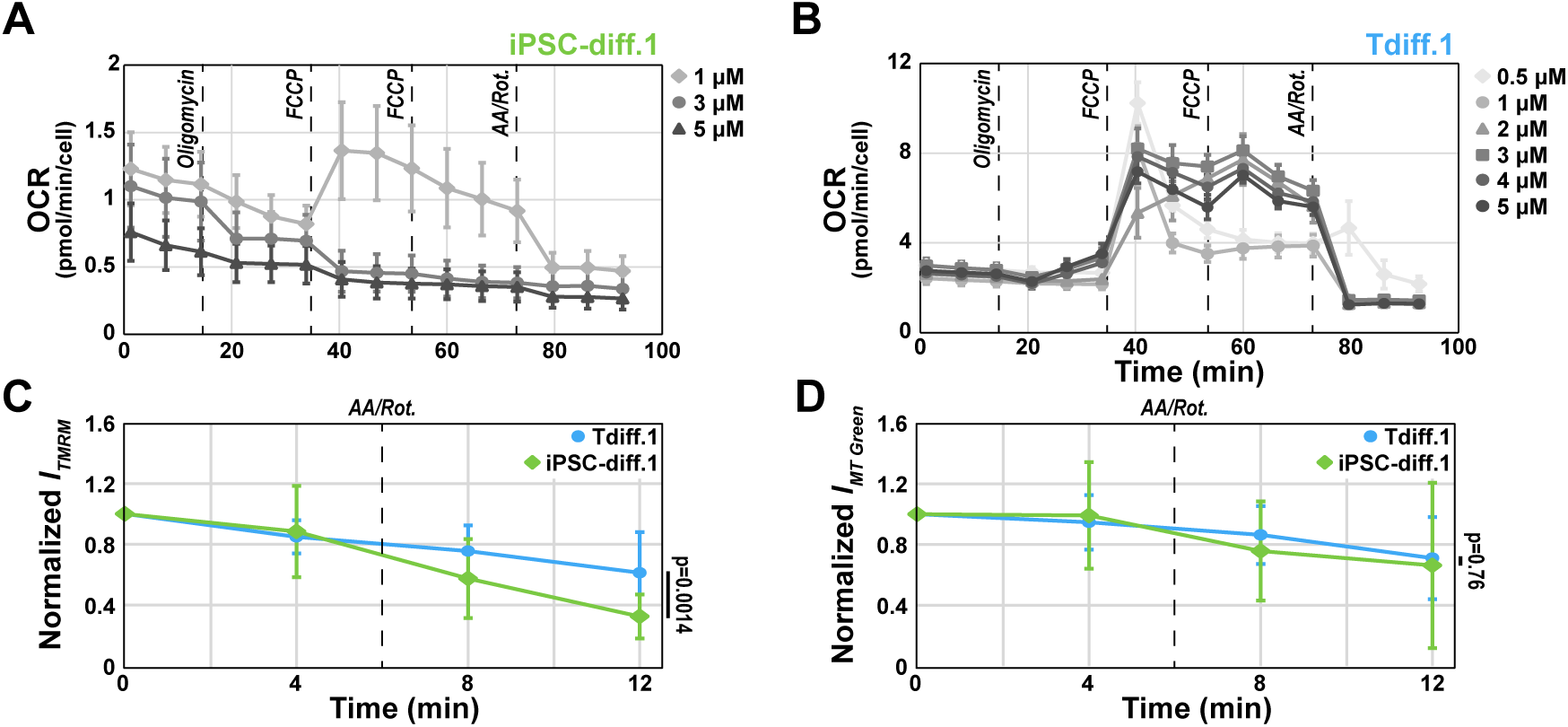
Increasing concentrations of mitochondrial stress molecules does not reduce proton leakage. **(A)** Mitochondrial stress test plots of oxygen consumption rate (OCR) over time in iPSC-diff.1 neurons. Mitochondrial stressors were added at the indicated time points at concentration listed in the legend. Error bars are standard deviation. **(B)** Same as (A) but for Tdiff.1 neurons. **(C)** Normalized intensity of TMRM over time during live-cell mitochondrial imaging. The dashed line denotes when antimycin A and rotenone were perfused into the cells. Error bars denote standard deviation, and statistics for the final timepoint were calculated using Welch’s t-test. **(D)** Same as (C) but for MitoTracker (MT) Green intensity.

**Figure S7:**
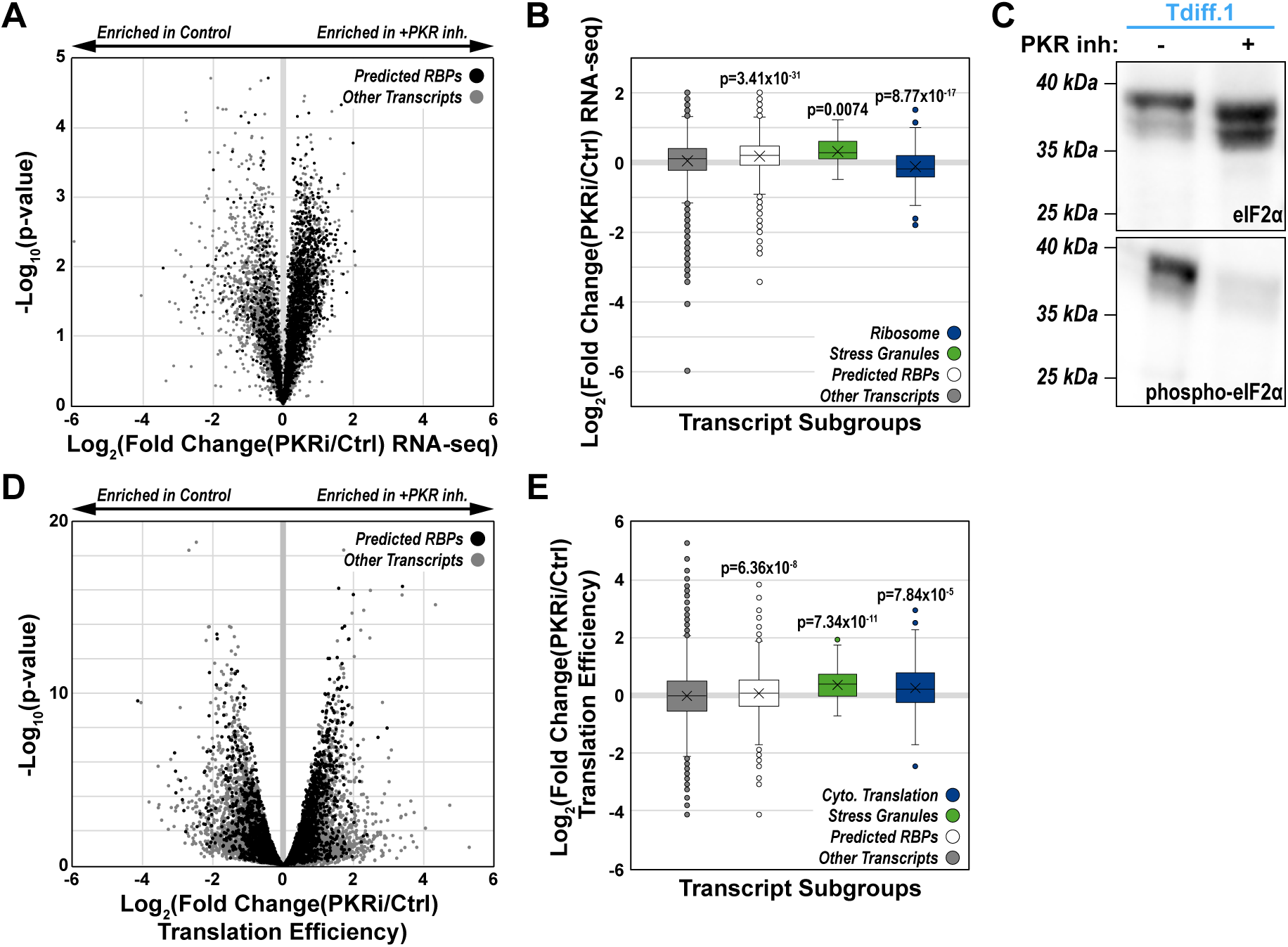
RNA-binding protein expression is rescued by PKR inhibition. **(A)** Volcano plot of RNA-seq expression of all detected transcripts in transdifferentiated neurons (Tdiff.1) with or without PKR inhibitor treatment (n=3 replicates). The p-values were calculated using a two-tailed Welch’s t-test. Predicted RNA-binding proteins (RBPs)^44^ are highlighted black. **(B)** Boxplot of the Log_2_(Fold Change) of all transcripts (gray), predicted RBPs (white), stress granule transcripts (green, GO:0010494), and ribosomal transcripts (blue, GO:0005840). From top to bottom, the box plot lines denote the 75^th^, 50^th^, and 25^th^ percentile of data values; error bars denote the range of non-outlier values, the “X” denotes the mean, and any indicated points are outliers. Statistics were calculated using a two-tailed Welch’s t-test for each treatment condition compared to the control. **(C)** Representative Western blot images of total and phospho-eIF2α in control and PKR inhibitor-treated Tdiff.1 neurons. **(D)** Same as (A) but for Riboseq data. **(E)** Same as (B) but for Riboseq data. In this plot, the ribosome subgroup was replaced by cytoplasmic translation (GO:0002181).

## Supplemental Tables

**Table S1:** List of top KEGG terms for the upregulated genes in transdifferentiated neurons.^a^.

| GO Term (Molecular Function) | Identifier | # of Genes | Fold Enrichment | Enrichment FDR |
| --- | --- | --- | --- | --- |
| Protein Digestion & Absorption | HSA04974 | 103 | 7.89 | 5.55x10<sup>-7</sup> |
| Legionellosis | HSA05134 | 57 | 7.13 | 9.78x10<sup>-4</sup> |
| Rheumatoid Arthritis | HSA05323 | 92 | 6.94 | 2.39x10<sup>-5</sup> |
| IL-17 Signaling Pathway | HSA04657 | 93 | 6.87 | 2.39x10<sup>-5</sup> |
| TNF Signaling Pathway | HSA04668 | 112 | 6.74 | 7.07x10<sup>-6</sup> |
| NF-κB Signaling Pathway | HSA04064 | 104 | 6.70 | 1.61x10<sup>-5</sup> |
| ECM-Receptor Interaction | HSA04512 | 88 | 5.94 | 5.28x10<sup>-4</sup> |
| Amoebiasis | HSA05146 | 102 | 5.69 | 3.59x10<sup>-4</sup> |
| Pertussis | HSA05133 | 76 | 5.35 | 0.00398 |
| Viral Protein Interaction with Cytokine and Cytokine Receptor | HSA04061 | 99 | 5.28 | 9.78x10<sup>-4</sup> |
| AGE-RAGE Signaling Pathway in Diabetic Complications | HSA04933 | 100 | 5.23 | 9.78x10<sup>-4</sup> |
| Alcoholic Liver Disease | HSA04936 | 141 | 4.12 | 0.00216 |

<sup>a</sup> Upregulated genes were determined by filtering for transcripts with an RPKM>1 and that were upregulated >4-fold in transdifferentiated neurons.

**Table S2:** List of top gene ontology terms for the significantly enriched dsRNA-interacting proteins.^a^.

| GO Term (Molecular Function) | Identifier | # of Genes | Fold Enrichment | Enrichment FDR |
| --- | --- | --- | --- | --- |
| Single-Stranded RNA Binding | GO:0003727 | 88 | 8.96 | 1.62x10<sup>-23</sup> |
| mRNA 3'UTR Binding | GO:0003730 | 102 | 8.59 | 2.66x10<sup>-25</sup> |
| Double-Stranded RNA Binding | GO:0003725 | 76 | 8.07 | 5.46x10<sup>-17</sup> |
| RNA Helicase Activity | GO:0003724 | 96 | 7.75 | 8.35x10<sup>-20</sup> |
| ATP-Dependent Activity Acting on RNA | GO:0008186 | 98 | 7.60 | 1.65x10<sup>-19</sup> |
| Translation Regulator Activity | GO:0045182 | 180 | 7.30 | 2.35x10<sup>-33</sup> |
| Helicase Activity | GO:0004386 | 183 | 6.82 | 5.52x10<sup>-30</sup> |
| mRNA Binding | GO:0003729 | 337 | 6.69 | 5.23x10<sup>-54</sup> |
| RNA Binding | GO:0003723 | 1852 | 6.36 | ~0<sup>b</sup> |
| Translation Regulator Activity Nucleic Acid Binding | GO:0090079 | 137 | 6.23 | 5.98x10<sup>-19</sup> |
| Ribonucleoprotein Complex Binding | GO:0043021 | 173 | 5.44 | 1.86x10<sup>-18</sup> |
| Catalytic Activity Acting on RNA | GO:0140098 | 455 | 4.43 | 3.56x10<sup>-32</sup> |
| GO Term (Cellular Component) | Identifier | # of Genes | Fold Enrichment | Enrichment FDR |
| Preribosome | GO:0030684 | 88 | 14.43 | 3.81x10<sup>-54</sup> |
| Cytoplasmic Stress Granule | GO:0010494 | 85 | 10.56 | 1.45x10<sup>-30</sup> |
| Polysome | GO:0005844 | 78 | 8.70 | 4.49x10<sup>-20</sup> |
| Cytosolic Ribosome | GO:0022626 | 122 | 7.36 | 3.05x10<sup>-23</sup> |
| Ribonucleoprotein Granule | GO:0035770 | 299 | 6.44 | 9.79x10<sup>-45</sup> |
| Ribonucleoprotein Complex | GO:1990904 | 807 | 6.43 | 1.73x10<sup>-126</sup> |
| Cytoplasmic Ribonucleoprotein Granule | GO:0036464 | 280 | 6.41 | 1.64x10<sup>-41</sup> |
| Nucleolus | GO:0005730 | 1126 | 5.46 | 9.92x10<sup>-132</sup> |
| Ribosome | GO:0005840 | 283 | 4.87 | 1.47x10<sup>-24</sup> |
| Nuclear Speck | GO:0016607 | 489 | 4.07 | 2.79x10<sup>-29</sup> |
| KEGG Term | Identifier | # of Genes | Fold Enrichment | Enrichment FDR |
| Ribosome Biogenesis in Eukaryotes | HSA03008 | 77 | 11.37 | 6.14x10<sup>-31</sup> |
| RNA Degradation | HSA03018 | 79 | 7.76 | 2.86x10<sup>-16</sup> |
| mRNA Surveillance Pathway | HSA03015 | 97 | 7.45 | 1.93x10<sup>-18</sup> |
| Spliceosome | HSA03040 | 131 | 4.85 | 5.35x10<sup>-11</sup> |
| Ribosome | HSA03010 | 134 | 4.74 | 7.85x10<sup>-11</sup> |
| Gap Junction | HSA04540 | 88 | 3.73 | 0.000201 |
| Pathogenic *E.coli* Infection | HSA05130 | 197 | 3.33 | 1.98x10<sup>-7</sup> |
| Nucleocytoplasmic Transport | HSA03013 | 108 | 3.24 | 0.00058 |
| Coronavirus Disease-COVID-19 | HSA05171 | 232 | 2.74 | 2.07x10<sup>-5</sup> |

<sup>a</sup> Enriched proteins were identified by filtering for any identified protein that was enriched >32-fold in J2/IgG.<sup>b</sup> The false discovery rate was too low to be calculated.

**Table S3:** Stress granule proteins in the dsRNA interactome of iPSC-diff.1 and Tdiff.1 neurons.

| Protein | iPSC-diff J2/IgG |  | Tdiff J2/IgG |  | FC-FC<sup>c</sup> |  | Protein | iPSC-diff J2/IgG |  | Tdiff J2/IgG |  | FC-FC<sup>c</sup> |
| --- | --- | --- | --- | --- | --- | --- | --- | --- | --- | --- | --- | --- |
|  | FC<sup>a</sup> | p<sup>b</sup> | FC<sup>a</sup> | p<sup>b</sup> |  |  |  | FC<sup>a</sup> | p<sup>b</sup> | FC<sup>a</sup> | p<sup>b</sup> |  |
| ATXN2 | 4.05 | 2.55 | 8.27 | 1.27 | 4.23 |  | MCRIP1 | 0.12 | 0.13 | 3.21 | 1.91 | 3.09 |
| ATXN2L | 2.43 | 2.09 | 6.24 | 1.15 | 3.81 |  | MCRIP2 | 0.79 | 0.80 | 4.46 | 1.19 | 3.67 |
| CAPRIN1 | 4.85 | 1.65 | 7.72 | 1.54 | 2.87 |  | MOV10 | 7.74 | 1.61 | 10.55 | 1.51 | 2.81 |
| CASC3 | 5.67 | 2.36 | 6.88 | 1.32 | 1.21 |  | NUFIP2 | 2.38 | 2.06 | 5.29 | 1.16 | 2.91 |
| CELF1 | 3.85 | 1.38 | 4.42 | 3.37 | 0.58 |  | NXF1 | 4.89 | 2.95 | 4.85 | 1.84 | -0.04 |
| CIRBP | 3.62 | 1.24 | 3.19 | 1.46 | -0.42 |  | PABPC1 | 4.98 | 1.65 | 6.52 | 2.18 | 1.53 |
| CSDE1 | 3.36 | 3.11 | 6.49 | 1.76 | 3.13 |  | PABPC3 | 0.96 | 0.73 | 0.88 | 0.55 | -0.07 |
| DDX1 | 2.89 | 2.54 | 4.69 | 1.82 | 1.81 |  | PABPC4 | 5.44 | 2.23 | 7.05 | 1.82 | 1.61 |
| DDX19A | 0.71 | 0.39 | 2.14 | 0.67 | 1.43 |  | PABPC5 | 7.10 | 1.92 | 7.79 | 1.34 | 0.69 |
| DDX3X | 2.67 | 3.40 | 4.19 | 1.30 | 1.52 |  | PKP1 | -3.70 | 0.58 | -1.94 | 0.42 | 1.76 |
| DDX6 | 2.63 | 3.40 | 5.51 | 1.57 | 2.88 |  | PQBP1 | 2.51 | 2.57 | 2.12 | 3.21 | -0.39 |
| DHX36 | 5.22 | 2.84 | 7.24 | 2.07 | 2.02 |  | PRRC2C | 3.64 | 2.47 | 7.34 | 1.19 | 3.70 |
| EIF2S1 | 1.25 | 0.90 | 2.77 | 1.58 | 1.52 |  | PUM1 | 2.72 | 1.66 | 7.31 | 1.47 | 4.59 |
| EIF3B | 1.15 | 1.21 | 2.80 | 2.75 | 1.65 |  | PUM2 | 3.71 | 2.46 | 7.06 | 1.44 | 3.35 |
| EIF4A1 | 0.49 | 0.39 | 1.38 | 1.20 | 0.89 |  | QKI | 4.02 | 1.67 | 3.67 | 1.36 | -0.35 |
| EIF4E | 0.80 | 0.87 | 3.64 | 1.12 | 2.84 |  | RBFOX1 | 1.51 | 1.81 | N.D.<sup>d</sup> | N.D.<sup>d</sup> | -1.51 |
| EIF4G1 | 2.15 | 2.45 | 4.63 | 1.33 | 2.47 |  | RBM4 | 3.35 | 3.11 | 4.49 | 2.67 | 1.14 |
| ELAVL1 | 4.84 | 2.01 | 4.45 | 1.60 | -0.39 |  | RBPMS | -0.30 | 0.15 | 1.06 | 0.49 | 1.36 |
| FMR1 | 5.02 | 2.76 | 7.13 | 1.40 | 2.11 |  | RC3H1 | 5.60 | 2.88 | 4.39 | 1.26 | -1.21 |
| FXR1 | 4.70 | 3.03 | 6.62 | 1.34 | 1.92 |  | RC3H2 | 6.99 | 1.99 | 6.64 | 1.73 | -0.35 |
| FXR2 | 5.13 | 4.99 | 6.85 | 1.39 | 1.71 |  | ROCK1 | 1.11 | 1.00 | 4.50 | 1.03 | 3.39 |
| G3BP1 | 3.98 | 1.94 | 5.06 | 1.61 | 1.08 |  | RPTOR | 2.06 | 1.03 | 4.76 | 1.65 | 2.70 |
| G3BP2 | 3.80 | 2.13 | 6.22 | 1.17 | 2.42 |  | SSB | 4.79 | 2.22 | 5.90 | 2.38 | 1.11 |
| GIGYF2 | 0.63 | 0.41 | 2.87 | 1.93 | 2.24 |  | STAU1 | 6.71 | 2.43 | 6.87 | 3.03 | 0.16 |
| HABP4 | 3.03 | 2.26 | 5.03 | 1.21 | 2.01 |  | STAU2 | 6.05 | 1.89 | 6.90 | 2.09 | 0.85 |
| HNRNPK | 2.81 | 1.68 | 4.11 | 1.99 | 1.30 |  | TARDBP | 1.33 | 1.38 | 2.11 | 1.78 | 0.78 |
| IGF2BP1 | 4.33 | 2.41 | 4.42 | 1.31 | 0.08 |  | TIA1 | 0.65 | 0.48 | 1.10 | 1.54 | 0.45 |
| IGF2BP2 | 4.71 | 2.33 | 6.18 | 1.37 | 1.46 |  | TIAL1 | 1.68 | 1.91 | 1.62 | 2.20 | -0.06 |
| IGF2BP3 | 4.15 | 1.85 | 6.00 | 1.70 | 1.85 |  | TRIM21 | 0.02 | 0.01 | 0.23 | 0.12 | 0.21 |
| KPNB1 | 1.54 | 1.11 | 2.13 | 1.85 | 0.59 |  | TRIM25 | 4.68 | 1.55 | 6.28 | 1.45 | 1.60 |
| L1RE1 | 7.13 | 1.41 | 0.37 | 0.37 | -6.76 |  | UBAP2L | 1.64 | 1.60 | 3.30 | 1.62 | 1.66 |
| LARP1 | 5.64 | 2.06 | 8.06 | 1.55 | 2.41 |  | VCP | 0.04 | 0.02 | -0.48 | 0.27 | -0.51 |
| LARP1B | 5.02 | 1.45 | 6.86 | 1.53 | 1.84 |  | YBX1 | 5.24 | 3.63 | 6.98 | 2.42 | 1.74 |
| LARP4 | 4.23 | 2.25 | 7.29 | 1.62 | 3.06 |  | YTHDF1 | 3.11 | 2.26 | 5.11 | 1.45 | 2.00 |
| LARP4B | 3.27 | 2.73 | 5.27 | 1.38 | 2.00 |  | YTHDF2 | 1.77 | 1.76 | 4.72 | 1.17 | 2.94 |
| LIN28A | 4.66 | 2.11 | N.D.<sup>d</sup> | N.D.<sup>d</sup> | -4.66 |  | YTHDF3 | 1.18 | 0.96 | 4.73 | 1.13 | 3.56 |
| LSM14A | 2.20 | 1.00 | 2.27 | 0.60 | 0.07 |  | ZNFX1 | 3.23 | 1.93 | 5.66 | 2.51 | 2.43 |
| MBNL1 | 0.59 | 0.37 | 2.96 | 5.57 | 2.37 |  |  |  |  |  |  |  |

<sup>a</sup> The fold change values are reported as Log<sub>2</sub>(Fold Change) for J2 divided by IgG.<sup>b</sup> The p-values are reported as -Log<sub>10</sub>(p-value) for J2 and IgG IP samples.<sup>c</sup> The FC-FC values are reported as the Log<sub>2</sub>(Fold Change-Transdiff.) – Log<sub>2</sub>(Fold Change-iPSC-diff.)<sup>d</sup> Samples that were not detected in both the IgG and J2 pulldown are reported as not detected (N.D.). Proteins in the stress granule GO term (GO:0010494) that were not detected in any sample are not included in the table.

**Table S4:** TPM values for MT-RNA transcripts in dsRNA-seq results of transdifferentiated neurons.

| Transcript | iPSC-diff.1 Average |  | Tdiff.1 Average |  | Tdiff.2 Average |  | Tdiff.5 Average |  |
| --- | --- | --- | --- | --- | --- | --- | --- | --- |
|  | Input | IP | Input | IP | Input | IP | Input | IP |
| MT-ATP6 | 1611.434 | 2778.534 | 7717.179 | 8834.358 | 7590.708 | 4739.07 | 5604.037 | 8953.228 |
| MT-ATP8 | 2692.787 | 7901.868 | 16758.04 | 25027.68 | 16851.79 | 10600.78 | 8648.36 | 20374.35 |
| MT-CO1 | 2356.635 | 8907.028 | 9078.815 | 25875.27 | 9158.473 | 12187.33 | 8390.537 | 16549.51 |
| MT-CO2 | 2005.457 | 4464.888 | 8711.44 | 12166.2 | 9083.018 | 7668.808 | 7816.798 | 16335.5 |
| MT-CO3 | 1905.458 | 3803.498 | 5938.421 | 6175.347 | 5959.622 | 4745.872 | 4030.27 | 6416.037 |
| MT-CYB | 969.9146 | 1955.343 | 5035.749 | 6140.703 | 5082.184 | 4094.406 | 3933.837 | 6236.038 |
| MT-ND1 | 989.9305 | 2287.721 | 6675.637 | 9247.771 | 6272.309 | 4969.002 | 3861.35 | 7143.078 |
| MT-ND2 | 815.9881 | 1729.104 | 5183.186 | 7715.796 | 5042.539 | 4190.268 | 3519.795 | 6045.663 |
| MT-ND3 | 912.18 | 1606.06 | 3593.184 | 3257.372 | 3817.819 | 2669.197 | 3312.418 | 3426.022 |
| MT-ND4 | 1709.918 | 3170.171 | 7888.15 | 10865.14 | 7675.773 | 5489.478 | 6242.64 | 9999.527 |
| MT-ND4L | 1512.188 | 2799.128 | 5499.53 | 8548.922 | 5465.356 | 4712.102 | 5666.507 | 7213.303 |
| MT-ND5 | 986.2402 | 2314.605 | 5569.058 | 10760.52 | 5301.667 | 5336.715 | 4255.738 | 7416.342 |
| MT-ND6 | 241.8952 | 965.5774 | 202.3068 | 586.5533 | 198.3626 | 714.4541 | 116.4804 | 142.1376 |
| MT-RNR1 | 63401.15 | 257271.1 | 224264.6 | 376505.9 | 248718.4 | 368459.4 | 208029.1 | 378704.9 |
| MT-RNR2 | 56018.35 | 158700.7 | 200393 | 333710.2 | 236473.9 | 268274.5 | 204909.6 | 322452 |
| *Total* | *138129.5* | *460655.3* | *512508.3* | *845417.8* | *572692* | *708851.4* | *478337.4* | *817407.6* |
| **Fraction** | **0.138** | **0.461** | **0.513** | **0.845** | **0.573** | **0.709** | **0.478** | **0.817** |

**Table S5:** TPM values for MT-RNA transcripts in dsRNA-seq results of human brain tissue.

| Transcript | Mid-Age Average |  | Old-Age Average |  |
| --- | --- | --- | --- | --- |
|  | Input | IP | Input | IP |
| MT-ATP6 | 2958.566 | 1978.895 | 2892.101 | 4735.655 |
| MT-ATP8 | 25709.22 | 17243.67 | 27037.4 | 50019.93 |
| MT-CO1 | 2498.982 | 2162.681 | 2027.186 | 6027.006 |
| MT-CO2 | 2268.815 | 2056.098 | 2248.69 | 5095.4 |
| MT-CO3 | 2517.094 | 1655.471 | 1835.696 | 3405.733 |
| MT-CYB | 1061.116 | 672.6314 | 797.8821 | 1499.749 |
| MT-ND1 | 1706.884 | 1705.334 | 1778 | 4352.476 |
| MT-ND2 | 1817.913 | 1417.659 | 2061.522 | 4424.016 |
| MT-ND3 | 1337.247 | 1575.902 | 1387.02 | 3670.049 |
| MT-ND4 | 2153.068 | 1446.481 | 1631.801 | 3391.48 |
| MT-ND4L | 3199.583 | 3149.6 | 2956.795 | 6493.641 |
| MT-ND5 | 968.2765 | 805.7627 | 683.2438 | 1732.255 |
| MT-ND6 | 70.97416 | 976.0064 | 56.40174 | 489.7184 |
| MT-RNR1 | 42906.28 | 69145.93 | 44806.83 | 176129.1 |
| MT-RNR2 | 58302.39 | 50384.37 | 58504.89 | 115349.6 |
| *Total* | *149476.4* | *156376.5* | *150705.5* | *386815.8* |
| **Fraction** | **0.149** | **0.156** | **0.150** | **0.387** |

**Table S6:** List of top gene ontology terms for significantly enriched G3BP1-interacting proteins following PKR inhibitor treatment in Tdiff.1 neurons.^a^.

| GO Term (Cellular Component) | Identifier | # of Genes | Fold Enrichment | Enrichment FDR |
| --- | --- | --- | --- | --- |
| Cytosolic Large Ribosomal Subunit | GO:0022625 | 57 | 33.13 | 2.95x10<sup>-20</sup> |
| Cytosolic Ribosome | GO:0022626 | 122 | 28.22 | 1.25x10<sup>-34</sup> |
| Small Ribosomal Subunit | GO:0015935 | 105 | 25.39 | 1.98x10<sup>-25</sup> |
| Cytosolic Small Ribosomal Subunit | GO:0022627 | 64 | 24.30 | 1.44x10<sup>-14</sup> |
| Ribosomal Subunit | GO:0044391 | 222 | 23.52 | 4.47x10<sup>-49</sup> |
| Large Ribosomal Subunit | GO:0015934 | 121 | 21.11 | 1.99x10<sup>-22</sup> |
| Polysome | GO:0005844 | 78 | 19.94 | 1.54x10<sup>-13</sup> |
| Ribosome | GO:0005840 | 283 | 18.45 | 5.87x10<sup>-44</sup> |
| Cytoplasmic Stress Granule | GO:0010494 | 85 | 18.29 | 5.11x10<sup>-13</sup> |
| Organellar Ribosome | GO:0000313 | 101 | 17.60 | 1.98x10<sup>-14</sup> |
| Mitochondrial Ribosome | GO:0005761 | 101 | 17.60 | 1.98x10<sup>-14</sup> |
| Ribonucleoprotein Complex | GO:1990904 | 807 | 13.08 | 1.09x10<sup>-79</sup> |
| KEGG Term | Identifier | # of Genes | Fold Enrichment | Enrichment FDR |
| Ribosome | HSA03010 | 134 | 29.01 | 1.36x10<sup>-39</sup> |
| Ribosome Biogenesis in Eukaryotes | HSA03008 | 77 | 14.43 | 5.11x10<sup>-08</sup> |
| Coronavirus disease-COVID-19 | HSA05171 | 232 | 14.36 | 2.85x10<sup>-24</sup> |
| Spliceosome | HSA03040 | 131 | 11.02 | 7.46x10<sup>-9</sup> |
| Nucleocytoplasmic Transport | HSA03013 | 108 | 7.20 | 0.0013 |
| mRNA Surveillance Pathway | HSA03015 | 97 | 5.73 | 0.030 |
| Amyotrophic Lateral Sclerosis | HSA05014 | 364 | 3.05 | 0.030 |

<sup>a</sup> Enriched proteins were identified by filtering for any identified protein that was enriched >4-fold in PKRinh./Control.

**Table S7:** List of cell lines used in this paper.

| Identifier | Age | Sex | Source | Identifier |
| --- | --- | --- | --- | --- |
| Tdiff.1<sup>a</sup> | 52 | F | Rhine *et al.*, 2025 | n/a |
| iPSC-diff.1<sup>a</sup> | 52 | F | Rhine *et al.*, 2025 | n/a |
| Fibroblast.1<sup>a</sup> | 52 | F | Rhine *et al.*, 2025 | n/a |
| Tdiff.2 | 63 | M | Rhine *et al.*, 2025 | n/a |
| Tdiff.4 | 85 | F | NIA; Rhine *et al.*, 2025 | AG13077 |
| Tdiff.5 | 43 | F | NIA | AG08692 |

<sup>a</sup> Tdiff.1 is isogenic to the iPSC-diff.1 and Fibroblast.1 lines used throughout the paper.
<sup>b</sup> The reprogramming step reset the age of the iPSC-derived neurons<sup>11</sup>.

**Table S8:** Aged human brain tissue used in this manuscript.

|  | ID# | Age | Gender |
| --- | --- | --- | --- |
| Mid-Age Cohort | Case 20<sup>a</sup> | 38 | M |
|  | Case 26 | 49 | M |
|  | Case 131 | 56 | M |
| Old-Age Cohort | Case 65 | 82 | M |
|  | Case 67<sup>a</sup> | 77 | M |
|  | Case 103 | 92 | F |

<sup>a</sup> The RNA quality scores were too low for Cases 20 and 67 to move forward with PKR CLIP and dsRNA-seq (Figure 3F-H, Table S5); these samples were not included in the cohorts for these experiments.

**Table S9:**
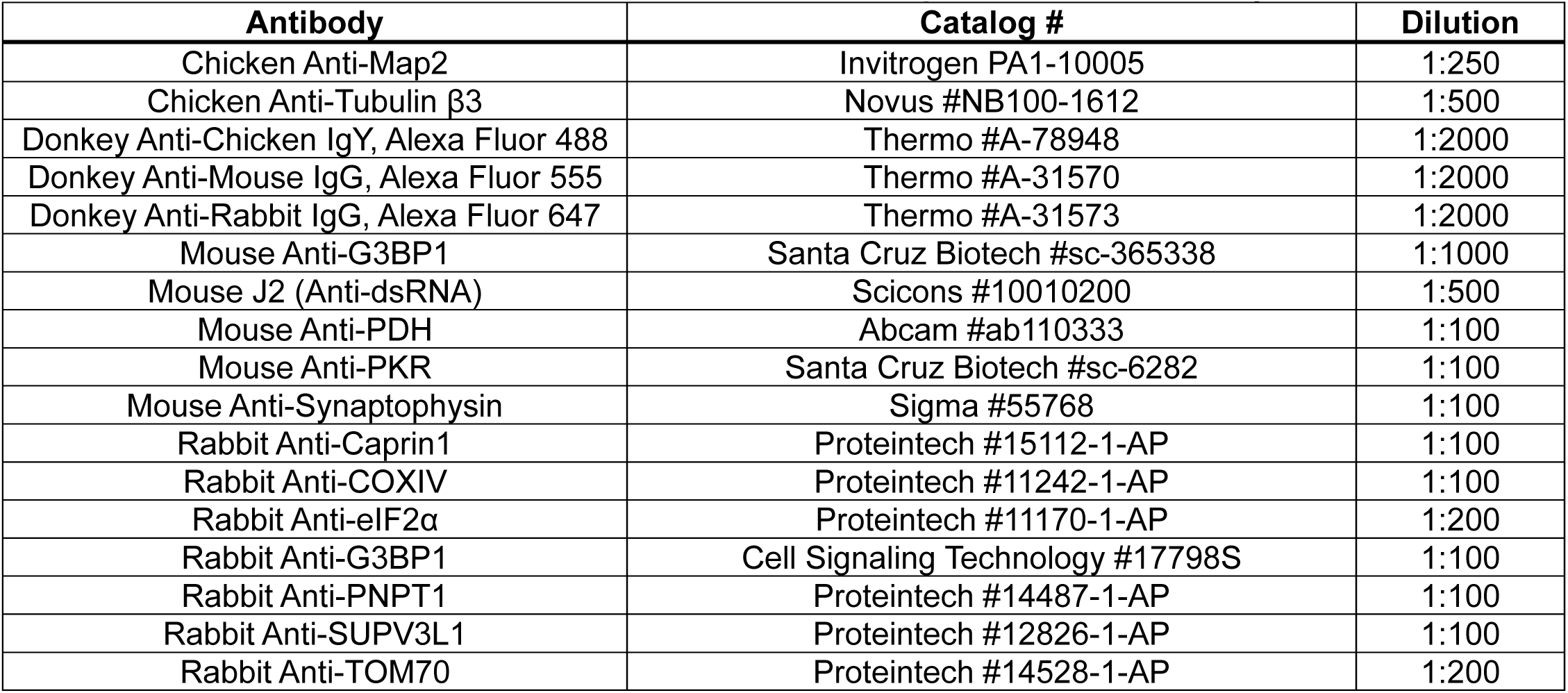
List of antibodies used for immunofluorescence experiments in this study.

